# Beyond Integration: SuperGLUE Facilitates Explainable Training Framework for Multi-modal Data Analysis

**DOI:** 10.1101/2024.11.19.624293

**Authors:** Tianyu Liu, Jia Zhao, Hongyu Zhao

**Affiliations:** Interdepartmental Program in Computational Biology & Bioinformatics, Yale University, New Haven, 06511, CT, USA; Department of Biostatistics, Yale University, New Haven, 06511, CT, USA

**Keywords:** Single-Cell Sequencing, Multi-Omic Data Analysis, Embeddings, Gene Regulatory Network Inference, Perturbation Analysis

## Abstract

Single-cell Multi-modal Data Integration has been an area of active research in recent years. However, it is difficult to unify the integration process of different omics in a pipeline, and evaluate the contributions of data integration. In this manuscript, we revisit the definition and contributions of multi-modal data integration, and propose a novel and scalable method based on probabilistic deep learning with an explainable framework powered by statistical modeling to extract meaningful information after data integration. Our proposed method is capable of integrating different types of omic and sensing data. It offers an approach to discovering important relationships among biological features or cell states. We demonstrate that our method outperforms other baseline models in preserving both local and global structures and perform a comprehensive analysis for mining structural relationships in complex biological systems, including inference of gene regulatory networks, extraction of significant biological linkages, and analysis of differentially regulatory relationships.

## 1 Introduction

Single-cell sequencing [1, 2] has matured into a robust technology that can collect multi-omic information from individual cells of various tissues [3, 4], including gene expression (RNA) profiles [1], DNA accessibility (ATAC) profiles [5], surface protein abundance (Protein) profiles [6], DNA methylation (Meth) profiles [7], spatial information [8], and others. Moreover, sequencing methods such as Patch-seq [9] can generate multi-sensing data, including Morphology (Morp), gene expression, and electrophysiology (Elec) at the single-cell level. Integrating information from multi-modal data, including multi-omic data and multi-sensing data, can provide a compressive description of cells’ functionality. Moreover, broad collaborations, such as Human Cell Atlas [10], and ROSMAP [11], are generating an exponentially increasing amount of atlas-level data facilitated by various technologies. Considering the different experimental settings of datasets, integrating single-cell datasets from different sources and modalities is an important step in utilizing large-scale data to explore biological systems [4].

With many integration methods developed for multi-modal data, it is challenging to improve the existing methods. Some benchmarking papers [12–14] discuss the advantages and disadvantages of various methods for different tasks with multi-modal information. It is difficult to develop a unified framework that addresses all the limitations of the existing methods. Moreover, beyond data integration, it is worth investigating what we can learn from multi-modal data integration (MDI) through joint modeling. We expect that MDI can handle tasks which cannot be addressed with single-omic datasets [15], and MDI can drive novel biological discoveries by analyzing the relation of different modalities [16, 17].

Most MDI approaches choose to project multi-modal data into a joint space, including TotalVI (RNA+Protein) [18], MultiVI (RNA+Protein+ATAC) [19], MultiMAP (arbitrary multi-modal data) [20], GLUE (arbitrary multi-modal data without spatial information) [16], UnitedNet [17] (arbitrary multi-modal data), and SpatialGLUE [21] (arbitrary spatially-resolved multi-modal data). These methods can learn a joint cell-level latent space to represent the cells from multi-modal data into a space for visualization and evaluation. GLUE and UnitedNet, in particular, also study the relation among features across different omic data. For example, GLUE utilizes prior knowledge to construct a peak-gene network and further constructs a data-specific cis-regulatory network by running a permutation test, thus it can offer information for the data-specific gene regulatory network (GRN) [22]. UnitedNet utilizes SHAP [23] to explain the relation of features across two omic data in the training process. However, these two methods have limitations. The permutation test used by GLUE is affected by the random seeds thus it can give different numbers and sets of relationships under different seeds. For UnitedNet, the SHAP values do not provide confidence or statistical significance, and the definition of important feature-feature relation lacks biological justification. Moreover, all of these methods do not consider the balance between the preservation of the global structure and the preservation of the local structure under the context of prior biological knowledge [24]. Therefore, there is a need for MDI methods with the ability to generate confident biological discoveries.

In this paper, we introduce a method for the integration of multi-modal data with considerations of integration performance, multi-modal-specific information extraction, and discovery of feature relationships. Our method, noted as Super Graph-linked Unified Embeddings (SuperGLUE), includes the advantages of its base model as well as additional features. SuperGLUE not only effectively integrates multi-modal data into a joint space, but also facilitates the discovery of both feature-feature relationship and feature-(cell) state relationship. Our method is guided by prior biological knowledge, which allows us to explore data-specific biological systems. Moreover, SuperGLUE is regularized by statistical methods, thus we can assess statistical significance for our discovered relationships based on graph perturbation. In conclusion, our method answers questions beyond data integration and demonstrates the ability of multi-modal data to reveal specific relationships in complex biological systems.

## 2 Results

### Overview of SuperGLUE

There are two main contributions of SuperGLUE. First, SuperGLUE extends graph-linked embeddings with databases for different omic data and classifiers for cell-type labels (as classifier-guidance training), thus SuperGLUE can integrate multi-modal data with paired or unpaired relationships. Moreover, we fuse the embeddings from classifiers and encoders by Kullback-Leibler (KL) divergence [25] (as information mixture learning), to ensure that the optimization direction is consistent. For spatial transcriptomic data, we further incorporate graph neural networks (GNNs) [26, 27] to encode the expression levels in niches. This aspect is included in both the pre-training step and the fine-tuning step. Second, SuperGLUE includes a novel test based on graph perturbation [28, 29] to discover feature-feature relationships by adding or removing certain edges of the biological prior graph. Combining this test with SHAP values to analyze the relationship between features and cell states, we propose a unified framework to explain the process of modeling biological systems. This aspect is included into the post-training step. We summarize the landscape of SuperGLUE in Figures 1 (a) and (b).

**Fig. 1.**
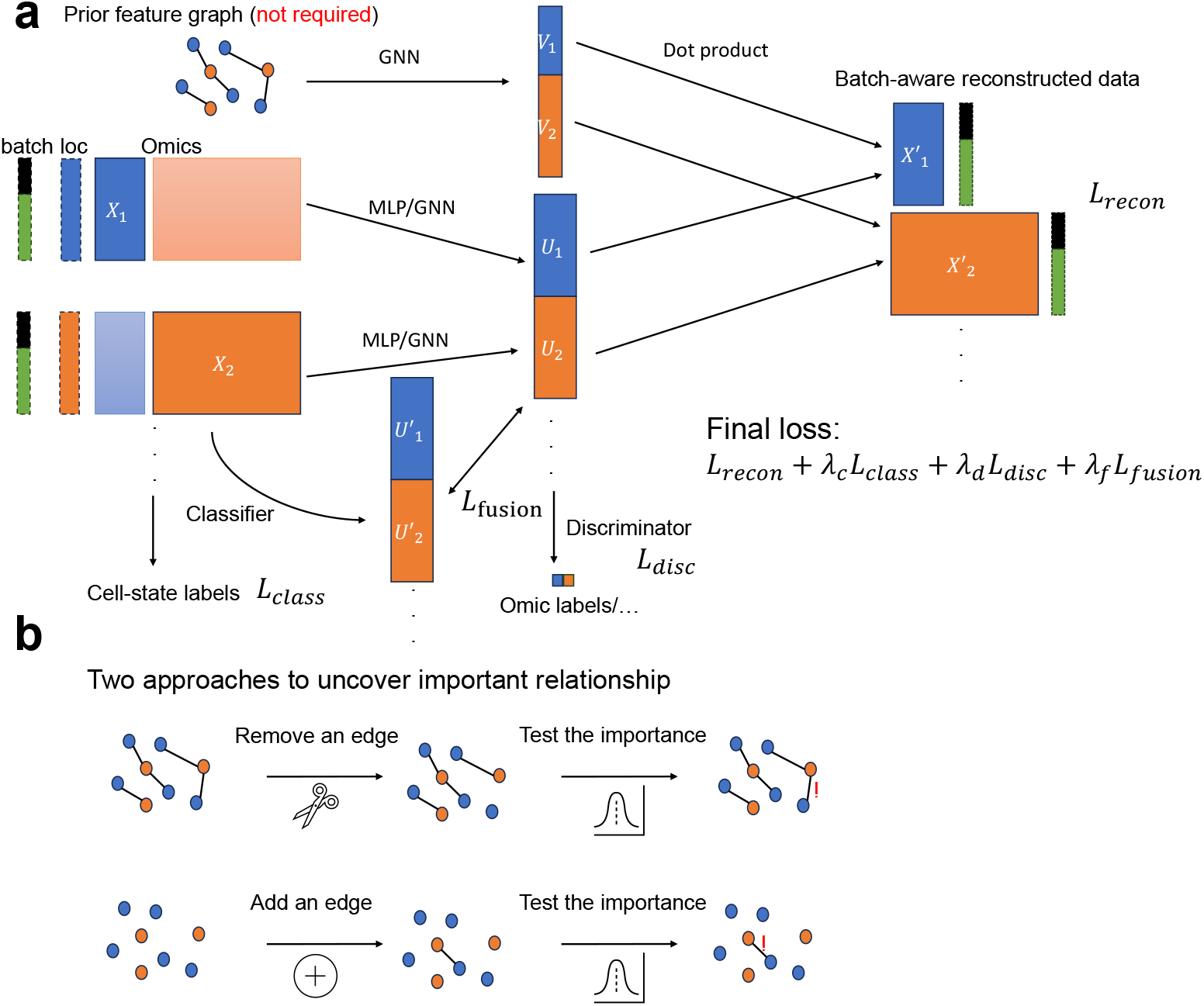
Overview of SuperGLUE. (a) The architecture of the SuperGLUE framework. We accept multi-modal datasets as well as biological prior graphs of features (optional) as inputs. We utilize MLP to encode non-spatial data and GNN to encode spatial data. GNN is always used to encode the information of the prior graph. The dot product mechanism is implemented to reconstruct the single-cell data. We have four loss components to balance multiple optimization targets, including reconstruction, cell-type classification (optional), omic label fusion, and latent embedding fusion. The optional metadata used for training are marked as dashed blocks. (b) The illustration of our statistical tests for uncovering important feature-feature relationships. We can either remove edges or add edges to test the difference of the node embeddings under two cases to compute test scores and significance levels.

### SuperGLUE enables the preservation of global structure

In the first section, we demonstrate the advantages of using SuperGLUE to integrate datasets in preserving the global structure of single-cell datasets compared with UnitedNet (another explainable framework) and GLUE. We first define the concepts of local structure and global structure in data visualization and data integration. The local structure [30] means the structure from a group of cells or samples, which is usually evaluated based on the k-nearest-neighbors. Instead, the global structure [30] considers the distance of different cell clusters. In data visualization, the balance of local structure and global structure has been argued for a long time [31, 32]. However, there has been little discussion on considering the trade-off. Moreover, most of the metrics used to evaluate data integration focus on local structure [33–35]. However, the global structure is also important, especially in describing the similarities and differences among cell states, which further support accurate cell-type classification and trajectory inference. Therefore, we propose a new metric for evaluating the global structure after data integration, known as PAGA [36, 37] similarity. The idea of PAGA similarity from [38] is to compute the global cell state information before and after integration to evaluate the performance of global structure preservation. We used the simulation datasets with two modalities (RNA and protein) from [17, 39] to perform experiments. Our visualization method is based on the top two principal components [40].

We display the global structure before integration in Figure 2 (a), and that after integration based on SuperGLUE In Figure 2 (b). There are no significant differences based on visualization. Furthermore, in Figure 2 (c), we compared SuperGLUE with UnitedNet based on PAGA similarity, where SuperGLUE and GLUE always outperformed UnitedNet under different input modalities. This is likely due to the fact that our method employs the structure of the Variational Auto-Encoder (VAE) [41] and reconstructs the distribution of the input data, thus mitigating the effect of noise in the single-cell data. UnitedNet utilizes an Auto-Encoder [42] structure and thus is not as good as our method at reconstructing the global structure.

**Fig. 2.**
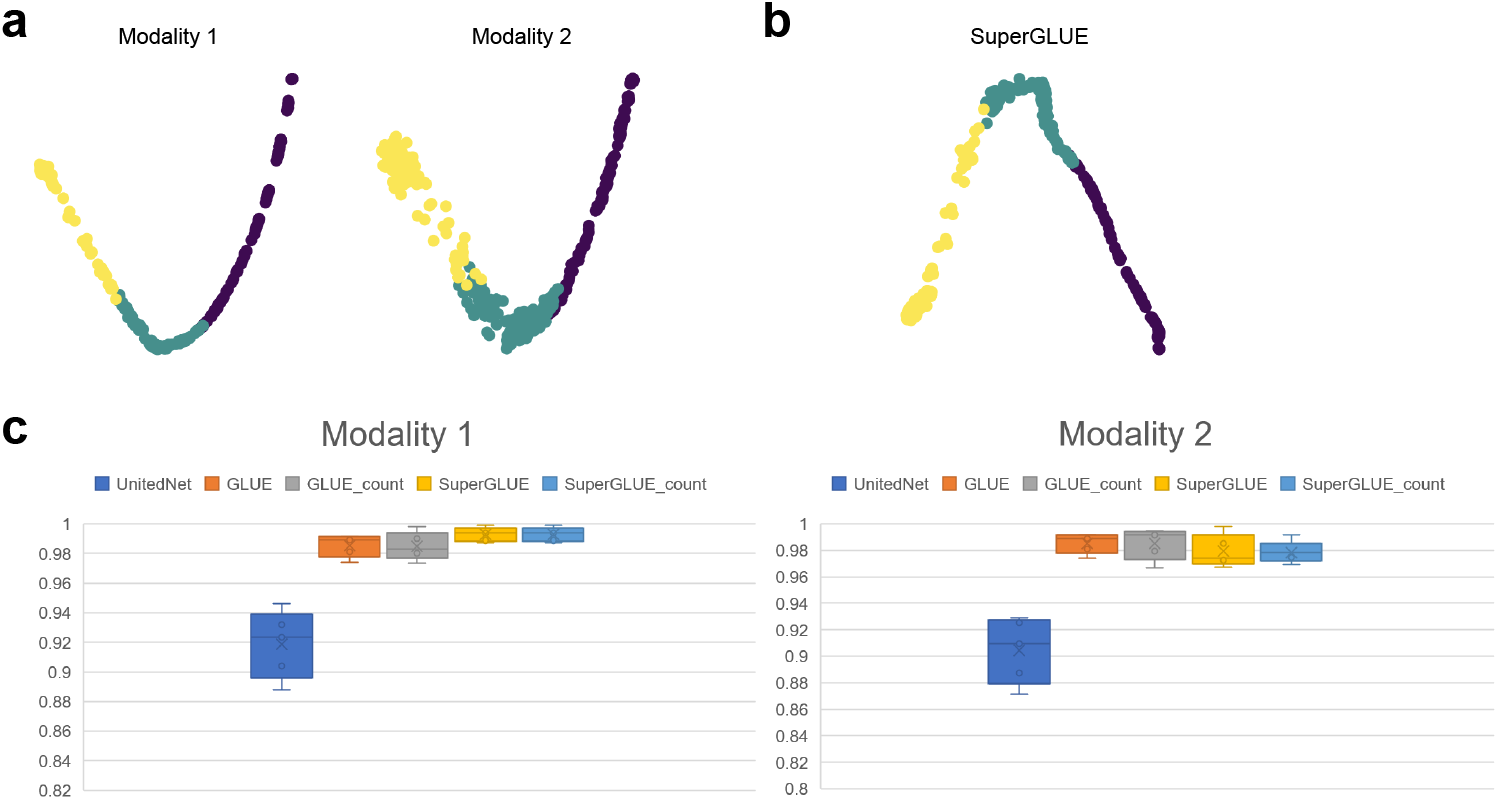
Comparisons of integration performances based on simulation datasets. (a) The PCA plots of the original multi-modal data. We illustrate these plots labeled by different types of modality. The color represents cell-type labels in the simulation. (b) The PCA plots of cell embeddings from SuperGLUE. The color represents cell-type labels in the simulation. (c) The boxplots of our metric to evaluate global structure preservation. We report the scores for five different simulation datasets. The observed structure is obtained from modality 1. (d) The boxplots of our metric to evaluate global structure preservation. We report the scores for five different simulation datasets. The observed structure is obtained from modality 2.

An interesting observation is that there is no major difference between SuperGLUE starting from principal components [40] and SuperGLUE starting from count data in preserving the global structure. However, considering computing efficiency, the former choice is better since its encoder has fewer parameters. Moreover, we investigated the relationship between the number of features of the first modality and PAGA similarity scores, shown in Extended Data Figure 1. This figure shows that a smaller number of features corresponds with a higher PAGA similarity score, suggesting that lower noise levels correspond to better global structure preservation.

These results show the superiority of SuperGLUE in preserving the global structure during data integration by comparing it with two leading methods, and we offered possible explanations for our discoveries.

### SuperGLUE outperforms other baselines in real-data analysis

In this section, we investigate the performance of SuperGLUE in integrating different types of multi-modal data by benchmarking different state-of-the-art (SOTA) methods. We evaluated the ability of both removing technical noise (known as *S*_*tech*_) and preserving biological information (known as *S*_*bio*_). For unpaired data integration, the technical noise comes from distinct omics platforms and batch effects, while for paired data integration, the technical noise only comes from batch effects. To evaluate *S*_*tech*_, we considered metrics from scIB [33], including tASW, iLISI, kBET, and Graph Connectivity (GC). To evaluate *S*_*bio*_, we also used metrics from scIB, including NMI, ARI, cASW, cLISI, and PAGA similarity, where the first four metrics are used to evaluate the preservation of the local structure and PAGA similarity is used to evaluate the preservation of the global structure. Furthermore, we computed the weighted average to derive the final score as *S*_*final*_ = 0.6*∗S*_*bio*_+0.4*∗S*_*tech*_. The same weights were used in scIB, and [43] demonstrated that the choices of weights do not significantly affect the final ranks of different methods. The baseline models will be introduced before the presentation of our experiments for different omics. All the methods were tuned to their best hyper-parameter settings for different datasets. Overall, we selected the following data integration scenarios: For the integration of two types of omic data, we considered scRNA+scATAC, scRNA+scProtein, scRNA+scMeth, and spRNA+spMorp. For the integration of three types of omic data, we considered scMorp+scRNA+scElec. Here, *sc* represents single-cell, and *sp* represents spatial. Our problem settings nearly covered all known multi-modal data types. We first present the results of bi-modality integration, as illustrated in Figure 3 (a).

**Fig. 3.**
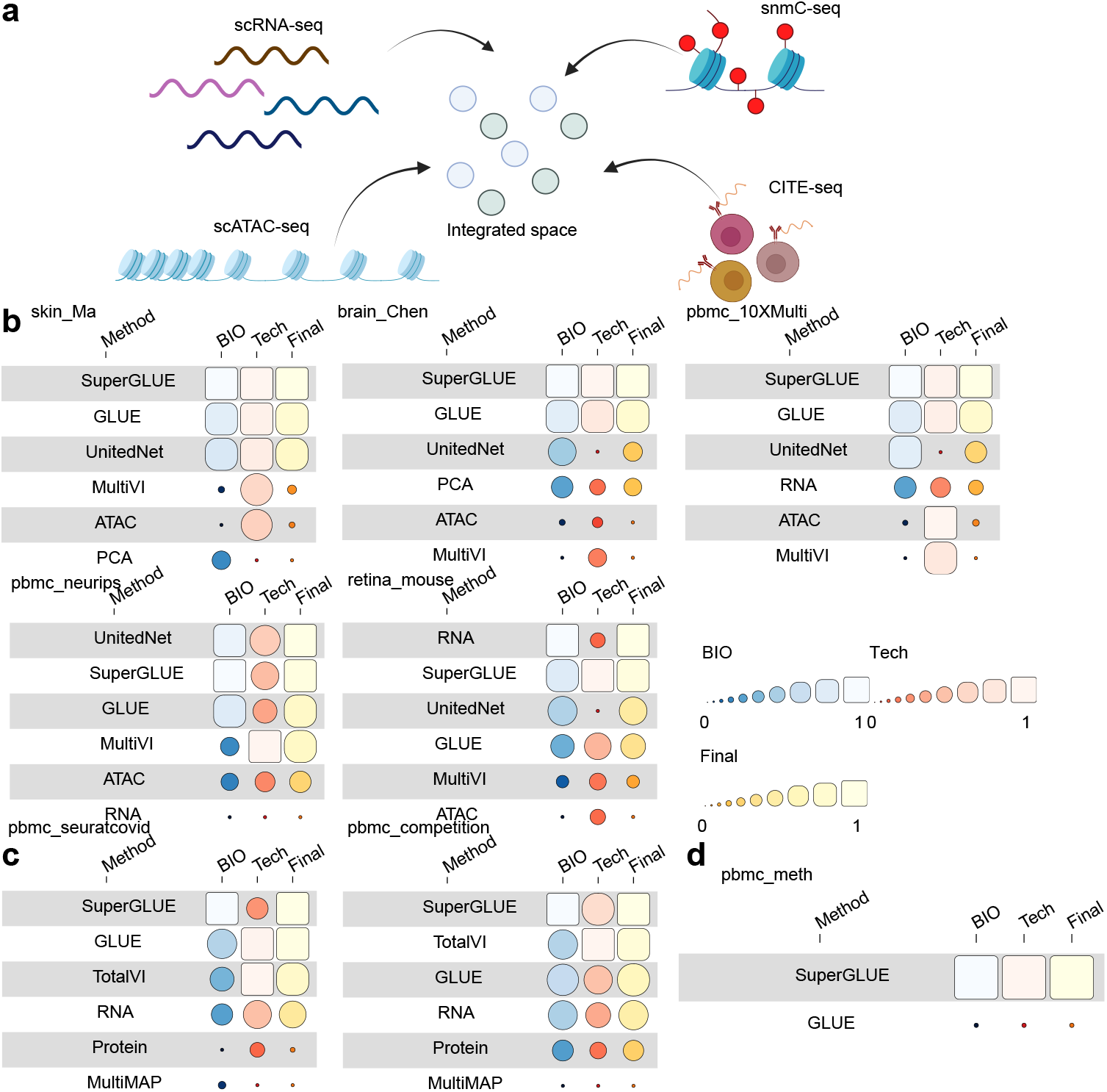
Comprehensive benchmarks for the bi-modality integration task. (a) The illustration of the bi-modality integration task. The idea is to integrate datasets from different omics into a joint latent space. (b) Benchmarking results of scRNA+scATAC integration. (c) Benchmarking results of scRNA+scProtein integration. (d) Benchmarking results of scRNA+scMeth integration.

Regarding the integration of scRNA+scATAC, we included UnitedNet, GLUE, and MultiVI as baseline models. We also considered using the original scRNA data after Principal Component Analysis (PCA) processing and the original scATAC data after Iterative Latent Semantic Indexing (LSI) processing [44, 45], as validation for the contribution of data integration. Here we considered five datasets for benchmarking, including skin_Ma [46], brain_Chen [47], pbmc_10XMulti (From 10X Genomics), pbmc_neurips [48], and retina mouse [49]. skin_Ma and pbmc_neurips contain real batch labels, while other datasets contain pseudo batch labels through random splitting. Here, all the datasets contain paired scRNA and scATAC information. According to Figure 3 (b), SuperGLUE showed a strong ability to integrate these two modalities by preserving the biological structure and reducing the technical effects. Our method ranked at the top in four out of five datasets and 2nd place in the Retina dataset. Moreover, SuperGLUE performed better than GLUE in every dataset, highlighting our implementation of classification guidance training and information mixture learning.

As for integrating scRNA+scProtein, we included (modified) GLUE, TotalVI, and MultiMAP as baseline models. UnitedNet has errors in integrating datasets from these two modalities. Since GLUE initially did not support the integration of scRNA and scProtein, we modified the original structure of GLUE. Here, we considered two datasets for benchmarking, including pbmc_seuratcovid [50, 51], and pbmc_competition [48]. According to Figure 3 (c), SuperGLUE still outperformed the other methods in integrating these two modalities, including GLUE. GLUE also has a shortcoming when interpreting the integration results of these two modalities: Since the number of proteins is much smaller than the number of genes, its permutation test fails as there is no significant protein-gene relationship. On the other hand, SuperGLUE overcomes this obstacle, which will be discussed in the section of model explainability.

Regarding the integration of scRNA+scMeth, we included GLUE as a baseline model. Here, scRNA and scMeth are unpaired data, and thus, the technical labels are the same as omic labels. We considered one dataset for benchmarking, denoted as pbmc meth [52, 53]. According to Figure 3 (d), SuperGLUE performed better than GLUE for both *S*_*tech*_ and *S*_*bio*_ scores. By considering all three cases, SuperGLUE always outperformed GLUE, and it achieved the best performances for most datasets.

Next, we investigate how SuperGLUE handles more complicated multi-modal datasets, including triple-omic data and multi-modal spatial data, which are illustrated in Figure 4 (a).

**Fig. 4.**
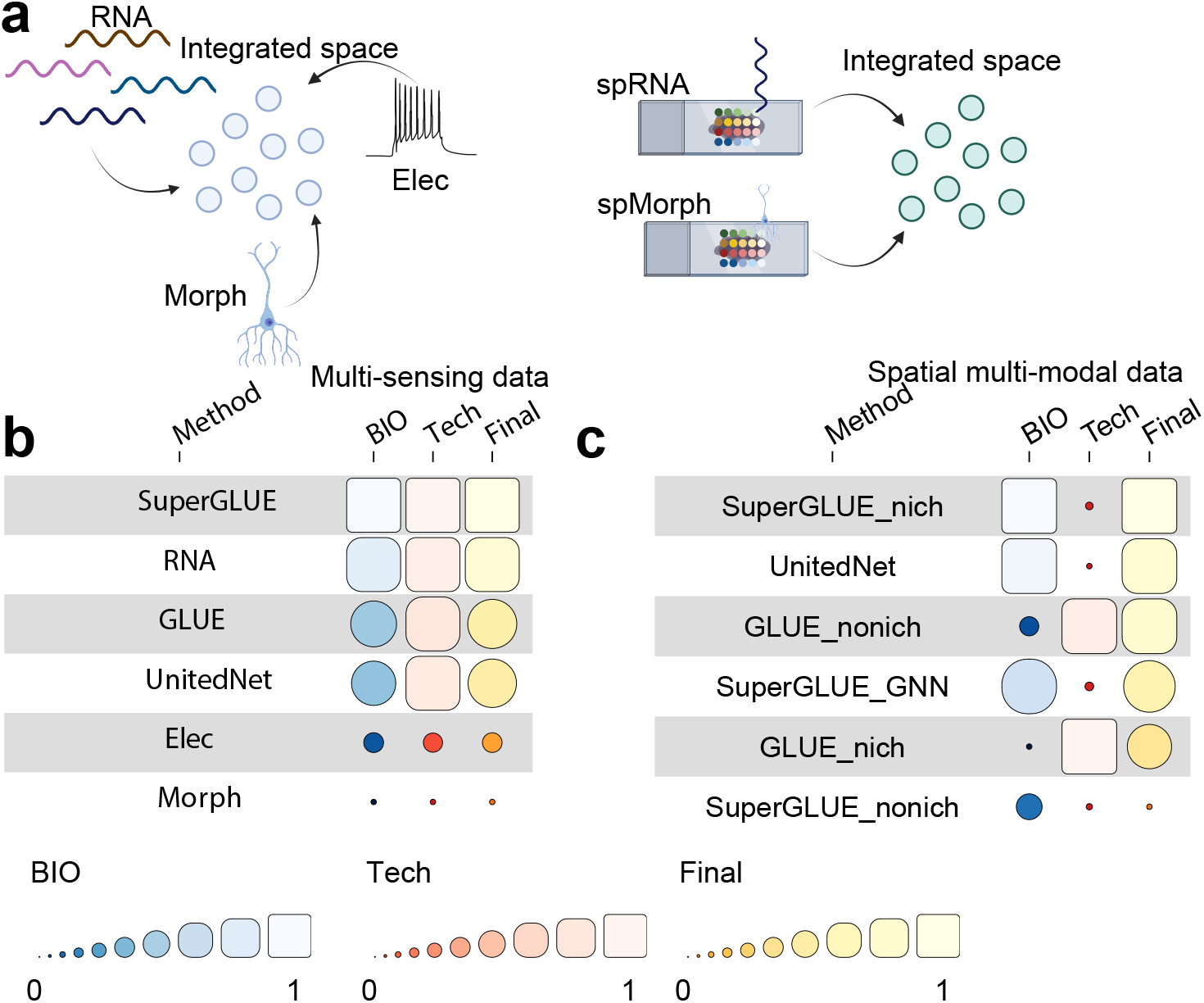
Comprehensive benchmarks for novel integration problems. (a) The illustration of two novel integration problems, which contain integration of the multi-sensing dataset and the spatially-resolved multi-modal dataset. (b) Benchmarking results for the integration of the multi-sensing dataset. (d) Benchmarking results for integration of the spatially-resolved multi-modal dataset.

Regarding the integration of scMorp+scRNA+scElec, we included UnitedNet and GLUE as baseline models. All the datasets came from the same cells through Patchseq [54]. We also considered using the original scMorph, scRNA, and scElec data after PCA processing in the benchmarking analysis. According to Figure 4 (b), SuperGLUE performed better than other baseline models in integrating the omic data from three different resources. Moreover, SuperGLUE outperformed GLUE again in this task and suggested that SuperGLUE’s improvement can be generalized. Meanwhile, we noticed that other baseline models had lower scores than the scRNA data representation. Therefore, SuperGLUE also has a better ability to utilize information from different omic data to formalize a better representation of cells. Since we do not have databases that link the features of these three omic datasets, we used these data as an example to illustrate the performance of feature discovery found by SuperGLUE’s test.

Regarding the integration of spRNA+spMorph, we included UnitedNet and GLUE as baseline models. The data are from [55], which contain mRNA information and morphological information as features. UnitedNet does not explicitly model the spatial information in the training process but utilized a weighted average approach to integrate the neighbors of the given spot, where the weight is inversely proportional to the distance between the spots. Such method is highlighted as “niche” transformation. Since GLUE does not support the integration of spatial data directly either, we included GLUE based on raw spatial data (GLUE nonich) as well as based on spatial data after “niche” transformation (GLUE nich) in our comparison. SuperGLUE GNN utilized GNN to incorporate the spatial information during the model training process, and we also considered SuperGLUE with and without ‘niche” transformation. According to Figure 4 (c), SuperGLUE nich ranked the 1st based on the final score. Meanwhile, it has performed relatively well in preserving biological structure. However, the GNN design of SuperGLUE outperformed the noniche design but performed worse than the niche design. These results demonstrated that integrating spatial multi-modal data required careful consideration of the approach to utilize geometry information.

### SuperGLUE is able to integrate atlas-level multi-modal datasets

In this section, we study the ability of SuperGLUE in integrating atlas-level multi-modal datasets. Our selected Fetal Atlas dataset [56] contains 377,624 cells, 1,154,464 peaks, and 36,601 genes. Our integrated results are shown in Extended Data Figures 2 (a) and (b). Based on Extended Data Figure 2 (a), SuperGLUE reduced the batch effect in the original multi-omic dataset, and the cell-type-specific information was preserved as well, shown in Extended Data Figure 2 (b). Therefore, SuperGLUE is capable of unifying different omic data into a space with a set of cell embeddings and has the potential to become a framework of a multi-omic foundation model [57]. Our method only requires one GPU with 16 GB memory (e.g., NVIDIA GTX 1080Ti) to integrate this atlas-level dataset, which is very efficient.

### SuperGLUE facilities explainability for multi-modal data analysis

Another important contribution of SuperGLUE is to formalize a well-defined framework for model explainability. Understanding the principles of factors in controlling and interacting with the biological process is an important task in computational biology [58, 59]. The deep learning method performs well in regression or classification problems [60, 61], but its black-box setting limits its power in addressing biological questions [62]. Therefore, we utilized two settings to discover different relationships inspired by [17, 28, 29]. To uncover the relationship between biological features and cell states, we relied on SHAP to explain their relationships and selected important biological features for each cell state. We proposed a novel statistical test based on graph perturbation to uncover the relationships among biological features from different omics. Details of our algorithms and tests are explained in the Methods section.

We first considered extracting important relationships between biological features and cell states, using important genes for cell states (also known as marker genes [63]) as examples. Based on the pbmc_10XMulti dataset, we visualize the results of SHAP in Figures 5 (a) for CD14 Mono and 5 (b) for D16 Mono. The results of all cell types after processing with SuperGLUE are shown in Figure 5 (c). The genes are ranked by the sum of absolute SHAP values. Higher SHAP values mean that higher values of such features lead to a higher probability of predicting the target cell type. We skipped the MT genes to avoid the confounders from aging [64], so genes DPYD and CTSS should correspond to each cell type. We further visualize the expression levels of these genes in Figure 5 (d), where DPYD has higher expression levels in CD14 Mono cells, while CTSS has higher expression levels in CD16 Mono cells. There is experimental support for these genes as markers [65, 66]. By including the top 20 (default number) genes ranked by the SHAP values for all cell types from the pbmc neurips dataset, we ran a Support Vector Classifier (SVC) with five-fold cross-validation to predict the cell types from pbmc_10XMulti dataset and compared the same results with all highly-variable genes. The results based on marker genes have better performances and significantly faster training speed, shown in Extended Data Figure 3. Therefore, our explainable learning framework can capture important biological features related to the given cell state, and the selected features can contribute to biological analysis.

**Fig. 5.**
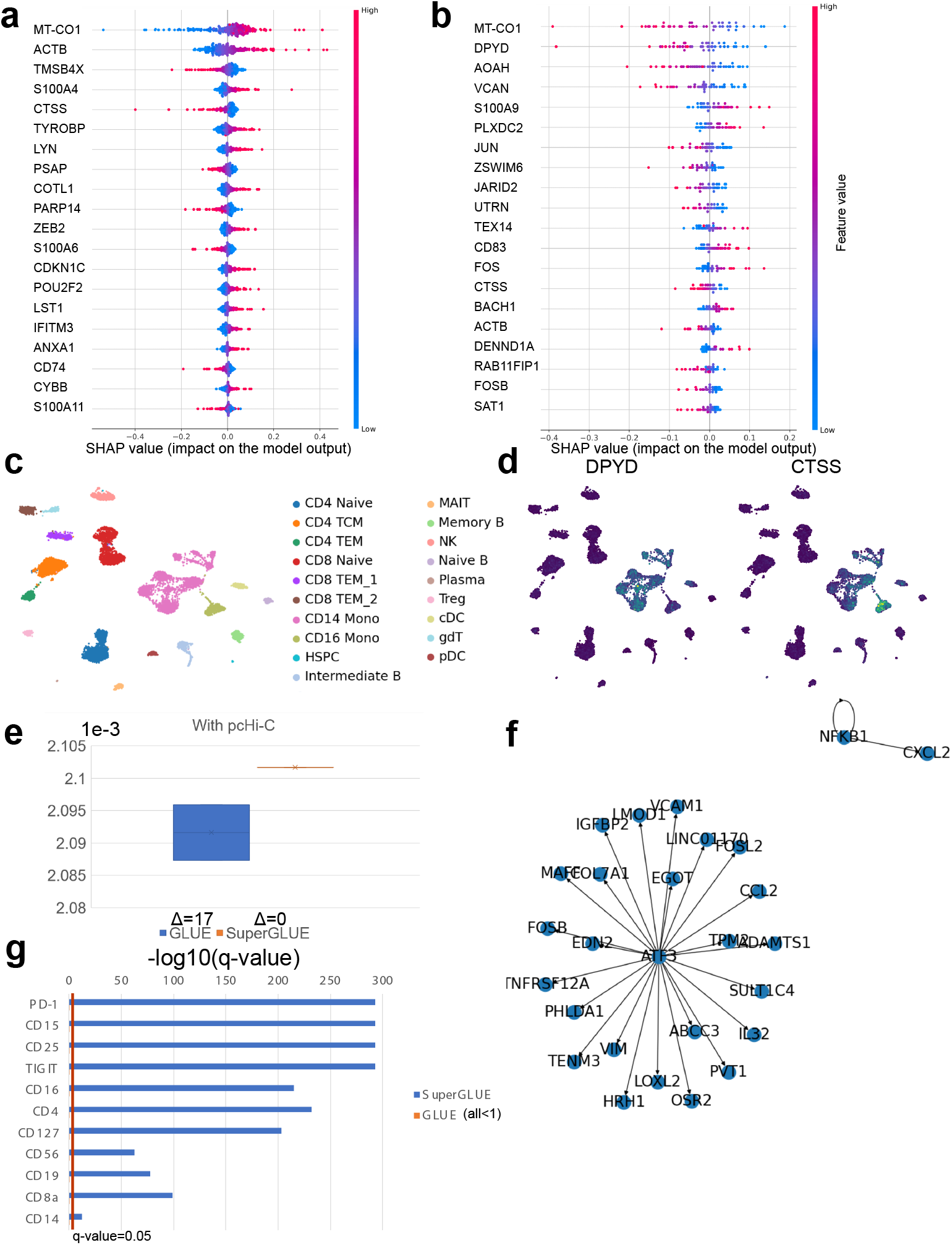
Explainable analyses from different aspects. (a) The scatter plot of SHAP values for the important genes inferred from CD14 Mono cells. (b) The scatter plot of SHAP values for the important genes inferred from CD16 Mono cells. (c) The UMAP plot of pbmc_10XMulti dataset colored by cell types. (d) The UMAP plot of pbmc_10XMulti dataset colored by selected gene expression levels, brighter color means a higher expression level. (e) The comparison of stability between SuperGLUE and GLUE for gene-peak relationship inference. The symbol Δ represents the difference between maximal number of relationships and minimal number of relationships under different seeds. (f) An example of GRN inferred based on gene embeddings from SuperGLUE based on the kidney dataset. (g) The important proteins uncovered by the protein embeddings from SuperGLUE based on the pbmc_3k5k dataset.

We then study the ability of our test in analyzing the relationship among biological features from different omics. Here, we utilized the dataset kidney muto to select significant gene-peak interactions. The “ground truth” information of such interactions can be extracted from either pcHi-C information [67] or eQTL information [68]. We set the threshold of adjusted p-value for significant interactions at 0.05, and compared the performances between GLUE with a permutation test and SuperGLUE with our novel test, shown in Figure 5 (e) based on pcHi-C and Extended Data Figure 4 based on eQTL. These two figures show that our test has lower variance and better overlap with pcHi-C information. The results of GLUE are affected by random seeds. Moreover, the permutation test and our test performed similarly by treating eQTL information as ground truth. Therefore, our test is not only stable (not affected by random seeds) but also has a higher overlap with known pcHi-C interactions. By utilizing the function of GRN inference in SCENIC/SCENIC+ [69, 70], we can also make inferences about GRNs by selected significant gene-peak interactions for this dataset. As shown in Figure 5 (f), and the inner loop of NFKB1 was also discovered in the Protein-Protein Interaction Network (PPIN) [71]. Therefore, SuperGLUE can also facilitate the discovery of novel and dataset-specific GRNs.

Another drawback of the permutation test is the requirement of a large sample size. For the pbmc 3k5k dataset, we had 11 proteins. By using our test from Super-GLUE, we discovered significant gene-protein interactions by setting a threshold of the adjusted p-value at 0.05, shown in Figure 5 (g). However, GLUE’s permutation test gives no significant results for gene-protein interactions. Proteins highlighted by our findings in significant interactions are further supported by biological experiments [72–81]. We also present the testing statistics of gene-protein interactions based on pbmc_seuratcovid and pbmc_competition in Extended Data Figures 5 (a) and (b), though we met running errors when benchmarking GLUE in these two datasets. Therefore, our statistical test not only realizes an alternative to the permutation test but also has a broader range of application settings.

All the cases we discussed above are the contributions of graph perturbation with edge removal. We utilized the Patch-seq dataset as an example to illustrate the contribution of graph perturbation with edge-adding settings, summarized in Appendix A. We also validated the correctness of the edge-adding settings by training a model without known edge information and discovering new gene-peak relationships in such a setting. We found that the overlap ratio between such two settings is larger than 95%. Therefore, these two settings are both reasonable and practical for different tasks.

### SuperGLUE is able to uncover specific biological networks and processes under perturbations

There are many computational methods for performing GRN inference [58, 82], and one important consideration of such research is the correctness of network inference, and thus, we need experiments to validate the inferred regulatory networks. These authors [83] generated Brachyury KO embryos by direct delivery of CRISPR/Cas9 as a ribonucleoprotein complex through electroporation method [84]. The perturbed dataset is named KO, while the control dataset is named WT. Such method can generate single-cell multi-modal datasets under perturbations, which highlighted a novel question for us to explore: How to consider and discover differentially expressed GRNs? Classical methods for GRN inference do not consider such a question. The cell-type distributions after integration are shown in Figures 6 (a) for the WT dataset and 6 (b) for the KO dataset. After perturbation, the composition of cell types changed. By utilizing SuperGLUE, we performed differentially expressed GRN inference for the KO and WT datasets. We first discovered dataset-specific gene-peak interactions by the statistical test, then performed SCENIC/SCENIC+ to infer the GRNs under different conditions. By analyzing these two sets of GRNs, we can identify the shared patterns and specific patterns for different conditions, which can be further explored to understand the biological process after perturbation. We visualized the GRN from the WT dataset in Figure 6 (c) and the GRN from KO dataset in Figure 6 (d). Qualitatively, we find that GRNs under normal conditions have a more connected set of nodes and that the transcript factors (TFs) in GRNs under the two conditions, as well as the regulatory relationships, are not identical. The GRN under the perturbed case has specific regulatory relationships dominated by TFs, including Foxq1, Hnf4a, klf12, and klf9. Experimental results also support the biological relationships between these TFs and Brachyury [58, 85–87]. Moreover, the TFs that were inferred to have differential regulatory relationships were different from the differentially expressed TFs inferred in the original paper of this dataset. Therefore, by analyzing the multi-modal data after integration, we can deconstruct biological processes at a deeper level, and the expression levels of genes are not as informative as embeddings of multi-modal features.

**Fig. 6.**
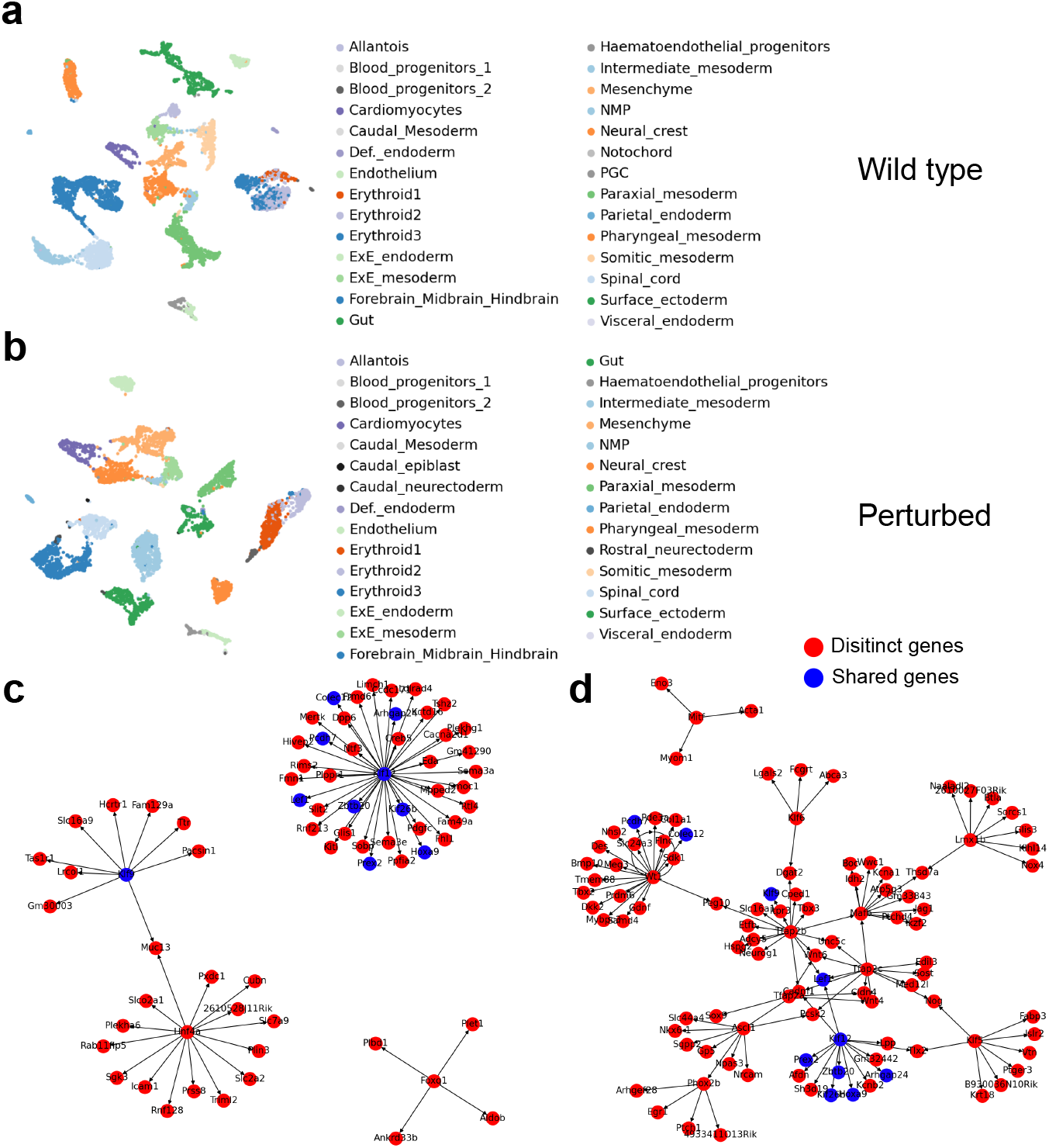
Analyses of differential regulatory relationships. (a) The UMAP plot of wild-type multiomic dataset based on cell embeddings from SuperGLUE colored by cell types. (b) The UMAP plot of a perturbed multi-omic dataset based on cell embeddings from SuperGLUE colored by cell types. (c) The GRN inferred based on the perturbed dataset. Nodes are colored by their overlapped condition. (d) The GRN inferred based on the wild-type dataset. Nodes are colored by their overlapped conditions.

Furthermore, we performed Gene Ontology Enrichment Analysis (GOEA) [88] for the gene sets in GRNs from both the WT dataset and KO dataset, shown in Extended Data Figures 6 (a) and (b). The threshold of adjusted p-values in this analysis is 0.05, and we visualized the top 10 pathways ranked by *−log*_10_(*adj p−values*). According to these figures, fewer pathways were significant after perturbation, and the top pathways showed differences between the two conditions. After perturbation, pathways involving neural activities and neurons are ranked in the top, highlighting the contributions of perturbations to uncover potentially important tissue-specific biological processes.

## 3 Discussion

The measurement of gene expression, along with other high-dimensional modalities from the same cells or different cells from the same tissues, poses exciting opportunities and challenges. Although many methods for MDI have been proposed to jointly analyze multi-modal information, there has been little research delving into the value of multi-modal data and/or biological questions that can be answered through these data.

In this paper, we have introduced a novel computational framework to perform data-driven biological discoveries and answer biological questions based on model outputs after data integration. We extend the graph-linked embeddings from multi-modal data with guided training and embedding fusion, and we also incorporate a GNN-based option to integrate spatial datasets. Our method is capable of various inputs and can be used to integrate different multi-modal tabular biomedical data at different single-cell level resolutions and large numbers of cells.

Our comprehensive benchmarking analysis demonstrates the better performance of SuperGLUE in integrating datasets from different modalities with good biological conservation and technical noise removal. We discuss metrics for evaluating the preservation of global biological structure and demonstrate that SuperGLUE is able to preserve both global and local biological structures. Our method also offers some statistical guarantee. To analyze the relationships between features and cell states, we use SHAP values to infer such relationships. Furthermore, we develop a statistical test based on graph perturbation to analyze the relationships among different biological features, which also has advantages in searching for important relationships in complex biological systems.

Although SuperGLUE has better performance over the existing methods in a number of aspects, there is room for improvement. For example, SuperGLUE offers statistics related to the explainability, the running time is long due to the computation of pairwise vector similarity. Moreover, We can also replace our current GNNs choices with more advanced ones or message-passing approaches to improve model performances. Moreover, we need to use the most recent, extensive, and reliable databases to guide our analysis. Overall, SuperGLUE provides new ideas for the application of multi-modal data analyses, and exploring novel applications after multi-modal data integration are essential for the development of computational biology.

## 4 Methods

### Problem statement

Consider a multi-modal sequencing dataset from 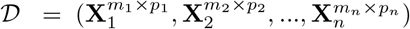, where the subscript represents the omic types and *m*_*i*_ represents the number of cells and *p*_*i*_ represents the number of features. For multi-modal information from the same cell, the number of cells for different modalities is the same, i.e., *m*_1_ = *m*_2_ = … = *m*_*n*_. Our target is to learn a model ℳ_*θ*_, which can integrate the multi-modal data into a joint space, that is:

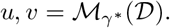

Here *u* represents the embeddings of cells and *v* represents the embeddings of biological features, in the joint space. Moreover, *γ*^*∗*^ represents the parameters of our models after training. *u* can be used for data visualization and benchmarking analysis, and *v* can be used for performing statistical tests to uncover significant relationships among biological features.

### Training process of SuperGLUE

SuperGLUE is a probabilistic deep learning framework, which is extended from GLUE. Therefore, we refer the Methods section in GLUE to help describe our method. The framework of SuperGLUE is based on Variational Auto-Encoder [41], which contains *n* encoders and *n* decoders for *n* omic data types. Considering the *k*^*th*^ omic dataset represented by 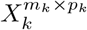, we can model this dataset by a low-dimensional variable in the joint space, which is denoted as cell embeddings *u*:

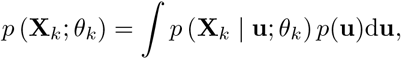

where *u* will be a variable shared by different omic datasets, and *p*(**u**) represents the prior distribution of the embeddings while *p* (**X**_*k*_ | **u**;*θ*_*k*_) represents the generative distributions to model the observed data. The generative distributions can be learned by the decoder component of our model.

Directly learning the posterior distribution *p* (**u** | **X**_*k*_; *ϕ*_*k*_) is difficult, so we introduced the variational posterior *q* (**u** | **X**_*k*_; *ϕ*_*k*_) to help us. We approximate the posterior distribution by the variational posterior. Therefore, the first loss function component of SuperGLUE is to minimize the negative value of the evidence lower bounds (ELBO):

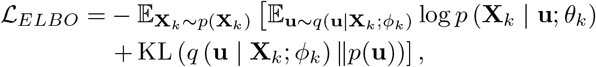

where *p* (**X**_*k*_) represents the data distribution of observed omic data, and *KL*() represents the KL divergence to minimize the probabilistic distance between *q* (**u** | **X**_*k*_; *ϕ*_*k*_) and *p*(**u**).

To incorporate the biological network information as prior information into our training process, we consider a graph as 𝒢 = (𝒱, ℰ), where 𝒱 represents the set of biological features (nodes) and *E* represents the known biological relationships (edges). The information of such graph can be extracted from databases. The biological feature embeddings *v* also come from the stackin𝒢 embeddings of 𝒱. In our example, all the edges have the same wights, but may have different signs under different biological contexts. For the relationships between genes and peaks, and those between genes and encoded proteins, the sign of edges is positive. For the relationships between genes and DNA methylation, the edges are negative.

Referring from our modelling process for omic data, we also treat the graph as an observed variable and build a VAE to learn the feature embeddings. By considering the information from the prior graph, we can rewrite the likelihood of SuperGLUE as:

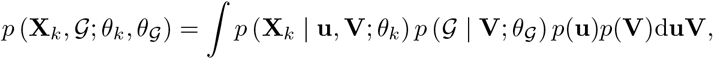

where *p* (𝒢 | **V**; *θ*_𝒢_) represents the generative distribution of biological prior graph, and *p*(**V**) represents the prior distribution of **V**. Therefore, we can learn the feature embeddings by constructing a similar ELBO. We also utilize the negative sampling approach [89] to improve the model performance according to (). The loss function is denoted as ℒ_𝒢_.

To reconstruct the omic dataset, our decoder is built on the inner product between the cell embeddings and feature embeddings. To initialize the cell embeddings, we can either use random vectors or embeddings from principal component analysis (PCA), which is depended the modes of SuperGLUE (whether starting from the data matrix in the expression space). To increase the capacity of our decoder, we choose to use the maximal likelihood method to optimize our ELBO, that is, we can model the input omic datasets under different distributions and fit their corresponding parameters. For example, if we intend to model the count-based scRNA-seq data and scATAC-seq data, we can use negative binomial (NB) distribution as follows:

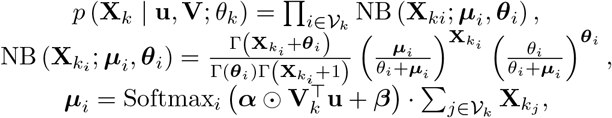

where *i, j* represent the indexes of features; *µ* represents the mean and *θ* represents the dispersion of NB distribution; *α* is a factor to learn the scale and *β* is a parameter to learn the bias; *Softmax*() represents the softmax function; and ⊙ represents the Hadamard product. To reduce the batch effect, we can model the parameters of our distributions for each batch, and treat the decoder as batch-specific.

To construct the encoder for cell embeddings **u**, we utilize multilayer perceptrons (MLPs) for datasets without spatial information and Graph Convolutional Networks (GCNs) for datasets with spatial information. To construct the encoder for feature embeddings **v**, we utilize GCNs for all the datasets.

Therefore, we have the first component of our loss function, which is denoted as *L*_*recon*_ = ℒ_*ELBO*_ + *λ*_𝒢_ℒ_𝒢_, where *L*_*recons*_ has been aggregated for all the omic datasets in the training process.

To align the embeddings from different omics, we utilize the adversarial training strategy [90] to fuse the information from different omics. Our idea is to introduce a discriminator *D*, which is trained to minimize the classifier for omic labels. That is:

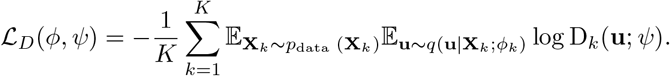

Here *ϕ* represents the parameters of our encoders and *ψ* represents the parameters of the discriminator. We also train the encoders to fool the discriminator, which can be treated as an asynchronous update training. To simplify our presentations, we unify the loss for adversarial training into one equation ℒ_*D*_. This is the second component of our loss function.

Inspired by the research about label-guidance learning, we also incorporate a classifier for cell types or cell states starting from the data matrix in the expression space, to formalize parts of the explainable framework, and improve model performance. We denote the classifier as *C*, and utilize the cross entropy loss, known as *L*_*class*_, to train the classifier. To enhance the data encoders’ performances, our omic-specific classifier is a 2-layer MLP and its first layer is used to generate cell embeddings **u**^**c**^. We also minimize the KL divergence between *q* (**u** | **X**_*k*_; *ϕ*_*k*_) and *q* (**u**^**c**^ | **X**_*k*_; *ϕ*_*k*_). The loss based on minimizing KL divergence is denoted as ℒ_*fusion*_. Therefore, we have the third and fourth loss components for our loss function.

Therefore, our final loss function is :

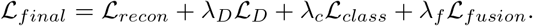

Here *λ*_*D*_, *λ*_*c*_, *andλ*_*f*_ are coefficients used for balancing the loss function components.

We note that we also inherit the weighted adversarial alignment and weighted embeddings averaging from GLUE, which are used to enhance the model’s performance for integrating atlas-level datasets. We tuned hyper-parameters of SuperGLUE for each dataset to achieve best performances, shown in Appendix B.

### Explainable framework for relationships between biological features and cell states

To uncover the relationships between biological features and cell states, we utilize *SHAP* () to extract the score of each feature corresponding to cell state *c*. The principle of SHAP comes from game theory, which tends to find the contributions of features towards predictions. *SHAP* () accepts a model and a dataset as input, and computes the SHAP score of each feature for the given label *c*, that is:

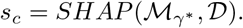

Here *s*_*c*_ is a vector whose length is equal to the number of biological features in *𝒟*. In our example, we analyzed the marker genes from different cell types by computing the SHAP values and ranked them based on single-cell multi-modal datasets. To verify the correctness of the selected marker genes, we consider both experimental validations and classification validations.

### Statistical test based on graph perturbation

After having the feature embeddings *v* from different omic datasets, we rely on graph perturbation to select important genes based on prior biological network (to find dataset-specific feature relationships) and to discover novel feature relationships (to find dataset-specific feature relationships without prior knowledge). Consider two omic datasets *o*_1_, *o*_2_, with corresponding feature embeddings 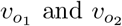, as well as a biological network 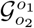. To generate the null distribution (where we assume the majority of the features are not correlated), we compute the cosine similarity for all the feature pairs based on the feature embeddings from these two omic datasets, that is:

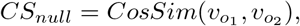

where *CS*_*null*_ is a vector of length 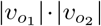. To select important feature relationships, we remove an edge from 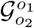, then calculate the new feature embeddings 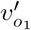 and 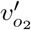 based on the perturbed graph (as 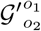). That is:

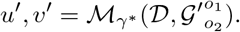

Therefore, we can compute the cosine similarity for all the feature pairs again, to generate the alternative distributions. That is:

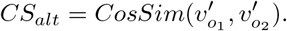

Finally, we utilize the Wilcoxon rank-sum test for these two distributions to determine if the selected edge is a significant edge. If we are interested in relationship discovery, we can add a new edge to 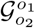, and compute the alternative distributions again. The significant feature relationships are selected with the following criteria: adj p-values*<*0.05. The adjusted p-values are computed based on False Discovery Rate (FDR) correction [91], same as scGLUE. The cases based on edge removal include cisregulatory inference and protein-gene interaction inference. The cases based on edge addition include joint analysis for the Patch-seq dataset.

### Gene regulatory network inference

The GRN inference is inspired by gene-peak regulatory relationship inference. Our pipeline is based on the ranking and pruning steps from SCENIC/SCENIC+. Our first step is to link peaks with TFs from ENCODE TF ChIP peaks [92]. Then for each gene, we compute the rank of their TF enrichment levels and rank the genes based on the p-values from the enrichment test. In this test, random matching between peaks and TFs is used to generate the null distribution. Finally, we produce the TF-gene networks and visualize them based on networkx [93]. We also validate the GRNs by measuring the overlap between inferred GRNs and a PPIN.

Regarding our analysis of differential GRNs, we perform the GRN inference step for the dataset under the controlled condition and the dataset under the perturbed condition. These datasets come from the same tissue and share cell-type distributions. We then produce the peak-gene regulatory relationships for each dataset followed by TF-gene networks. We compute the overlapped and specific parts for these two GRNs, and perform literature review to validate these findings.

### Explanations of evaluation metrics

The first set of metrics is for evaluating the performance of multi-modal data integration. Our settings are based on scIB [33] and PAGA [36]. We compute ASW_*tech*_, Graph Connectivity, kBET, and iLISI, and then average the scores from these metrics to generate *S*_*tech*_. We compute ASW_*label*_, NMI, ARI, cLISI, and PAGA similarity, and then average the scores from these metrics to generate *S*_*bio*_. Details of these metrics, referred from [33, 43], are introduced below:

1. PAGA Similarity: PAGA Similarity is a score to evaluate the conservation of global structure, under the biological context. The idea is to compute the connectivity scores as well as PAGA scores based on cell types before integration and after integration, and then we utilize the Gaussian kernel to evaluate the similarity of these two PAGA scores. The value of PAGA Similarity is between 0 and 1, and higher PAGA Similarity means better performances. For the simulation dataset, we consider evaluations based on the PAGA scores from two omics before integration. For real datasets, considering about the noisy level of multi-modal dataset, our PAGA score before integration is always computed based on scRNA-seq dataset.
2. Normalized Mutual Information (NMI): NMI is a score to evaluate the performance of biological information conservation. We compute this score based on the mutual information between the optimal Leiden clusters and the known cell-type labels and then take the normalization. The value of NMI is between 0 and 1, and higher NMI means better performance.
3. Adjusted Rand Index (ARI): ARI is a score to evaluate the performance of biological information conservation. ARI is used to evaluate the agreement between optimal Louvain clusters and cell-type labels. The value of ARI is between 0 and 1, and higher ARI means better performance.
4. Average Silhouette Width (ASW): We have cell type ASW (*ASW*_*cell*_) and technical ASW (*ASW*_*tech*_) for this metric. For one cell, ASW calculates the ratio between the inner cluster distance and the intra cluster distance for this cell. Therefore, higher *ASW*_*cell*_ means better biological information conservation and lower *ASW*_*tech*_ means better technical noise removal. To make them consistent, for *ASW*_*cell*_, we take the normalization, that is:

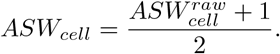

Similarly, for *ASW*_*tech*_, we take the inverse value of the normalized result, that is:

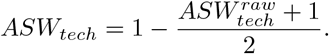

Both of metrics are in (0,1), and a higher score means better model performance.
5. Local Inverse Simpson’s Index (LISI): LISI is a metric to evaluate whether datasets are well-mixed under batch labels (*iLISI*) or can be discerned with different cell types *cLISI*. We first compute the k-nearest-neighbor list of one cell, and count the the number of cells that can be extracted from the neighbors before one label was observed twice. Furthermore, we take the normalization for *iLISI* with *B* batches, that is:

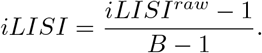

Similarly, for *cLISI* with *C* cell types, we take the inverse value of the normalized result, that is:

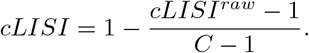

Both of metrics are in (0,1), and a higher score means better model performance.
6. Graph Connectivity (GC): GC measures the connectivity of cells in different cell types. If the batch effect is substantially removed, the connectivity of cells of the same cell type from different batches will have a higher connectivity score based on the k-NN neighbor graph. Therefore, we can compute the GC score for each cell type and take the average. GC score is in (0,1) and higher means better technical noise removal.
7. kBET: The kBET algorithm is used to determine if the label composition of the k-nearest-neighbors of a cell is similar to the expected label composition. For the batch label mixture of cells in the same cell type, the proportion of cells from different batches for the neighbors of one cell should match the global level distribution. The value of the kBET score is between 0 and 1, and higher score means better technical noise removal performance.

The second set of metrics is for evaluating the performance of cis-regulatory inference. Our ground truth information comes from pcHi-C and eQTL. We compute the Jaccard similarity between gene-peak relationships after inference and gene-peak relationships from ground truth databases as the metric to evaluate the performances of different methods for this task. The Jaccard similarity (JS) for two sets of gene-peak relationships *S*_1_, *S*_2_ is defined as:

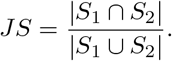

Here | | is used to compute the length of the given set. The value of the JS score is between 0 and 1, and higher score means better performance.

The third set of metrics is for evaluating the performance of GRN inference. Our ground truth information comes from PPIN. We utilize JS for two sets of TF-gene sets to evaluate the performance of models for this task. The JS score *∈* (0, 1) and higher score means better performance.

### Explanations of baselines

In this section, we summarize the major contributions of our baselines for multi-modal data integration. These methods are ranked in alphabetic order.

- **GLUE**: GLUE [16] is a model based on VAE and GNNs to integrate multi-modal datasets into a joint space, and it utilize a GNN to encode biological features into another space. The feature embeddinigs can be used to infer GRNs. The permutation test proposed by GLUE to select important relationships across biological features are not stable and practical for different types of single-cell multi-modal dataset, and it cannot be directly used to integrate spatial multi-modal datasets. SuperGLUE is an improved version of GLUE.
- **MultiMAP**: MultiMAP [20] is a manifold-learning-based method to integrate multi-modal datasets into a joint space. MultiMAP is capable of integrating different types of multi-modal datasets. However, it cannot integrate spatial multi-modal datasets, and does not perform well for different types of omic datasets.
- **MultiVI**: MultiVI [19] is based on VAE to integrate paired scRNA and scATAC datasets into a joint space. Its final embeddings are obtained by averaging the embeddings from different omics. Therefore, MultiVI has a limited range of applications.
- **TotalVI**: TotalVI [18] is based on VAE to integrate paired scRNA and scProtein datasets into a joint space. Its final embeddings are obtained by averaging the embeddings from different omics. Therefore, TotalVI has limited applications.
- **UnitedNet**: UnitedNet [17] is based on AE and a multi-task learning framework to integrate multi-modal datasets into a joint space. UnitedNet is capable of explaining the contributions of biological features as well as selecting important feature-feature relationships by SHAP. However, it has running errors based on SHAP and cannot offer statistical significance to support the discovered relationships. UnitedNet is also not good at preserving the global structure of biological datasets after integration.

Among these methods, only SuperGLUE, GLUE and UnitedNet can integrate datasets with three or more omic data types.

## Data availability

We did not generate new datasets in this project, and summarized the information of different datasets in Supplementary file 1.

## Reproductivity and Code availability

We used the resources from the Yale High Performance Center (Yale HPC) to conduct all of the experiments. Our maximal running time for each experiment was 24 hours. To train our model, we utilized one A5000 GPU with maximal RAM as 40 GB. To analyze our results, we utilized one CPU with maximal RAM as 150 GB. Our codes are released based on this link: https://github.com/HelloWorldLTY/SuperGLUE. The license is MIT license.

## 5 Acknowledgements

We appreciate the comments, feedback, and model explanations from Zhijie Cao, Gefei Wang. We also appreciate the PPIN dataset shared by Michelle M. Li.

## 6 Author contributions

T.L. designed the study with J.Z. T.L. ran all the experiments. T.L., J.Z. and H.Z. wrote the manuscript. H.Z. supervised this project.

## A Discovering novel relationships by graph perturbation

In this section, we discuss the contributions of our graph-perturbation-based test to select significant feature relationships. Based on ([17, 94, 95]), we integrate the multisensing dataset and select genes differentially expressed for Pvalb neurons. We ranked all the relationships based on p-values from smallest to largest and visualize the top 20 relationships in Extended Data Figures 7 (a), (b), and (c) for gene-morphology, gene-electrophysiology and morphology-electrophysiology, respectively. The genes Etv1, Syt17, Myo5b and Mtng1 all have strong linkage with the development or functionalities of neuron cells [96–99], and thus the linked morphology or electrophysiology also plays an important role in these cells. For example, bifurcation [100] is an important pattern in neuronal networks. Meanwhile, in our analysis, bifurcation relates with both gene expression levels and electrical signal measurements of cells.

## B Hyper-parameter tuning for SuperGLUE

In this section, we report the process to determine best hyper-parameter settings by using the 10X Multiome dataset as an example. In total, we consider adjusting *λ*_*c*_, *λ*_*d*_ and *λ*_*f*_. These parameters are hard to learn automatically because we do not have distribution assumption for every loss function component. The range for *λ*_*c*_ is [0.1,100], and the range for *λ*_*d*_ and *λ*_*f*_ is [0.01, 10]. Extremely large *λ*_*d*_ and *λ*_*f*_ will lead to model collapse. The *S*_*final*_ scores under different hyper-parameters are shown in Extended Data Figures 8 (a)-(c), which support us to select *λ*_*c*_ = 1, *λ*_*d*_ = 1 and *λ*_*f*_ = 100 in our final implementation.

## C Supplementary figures

**Extended Data Fig. 1.**
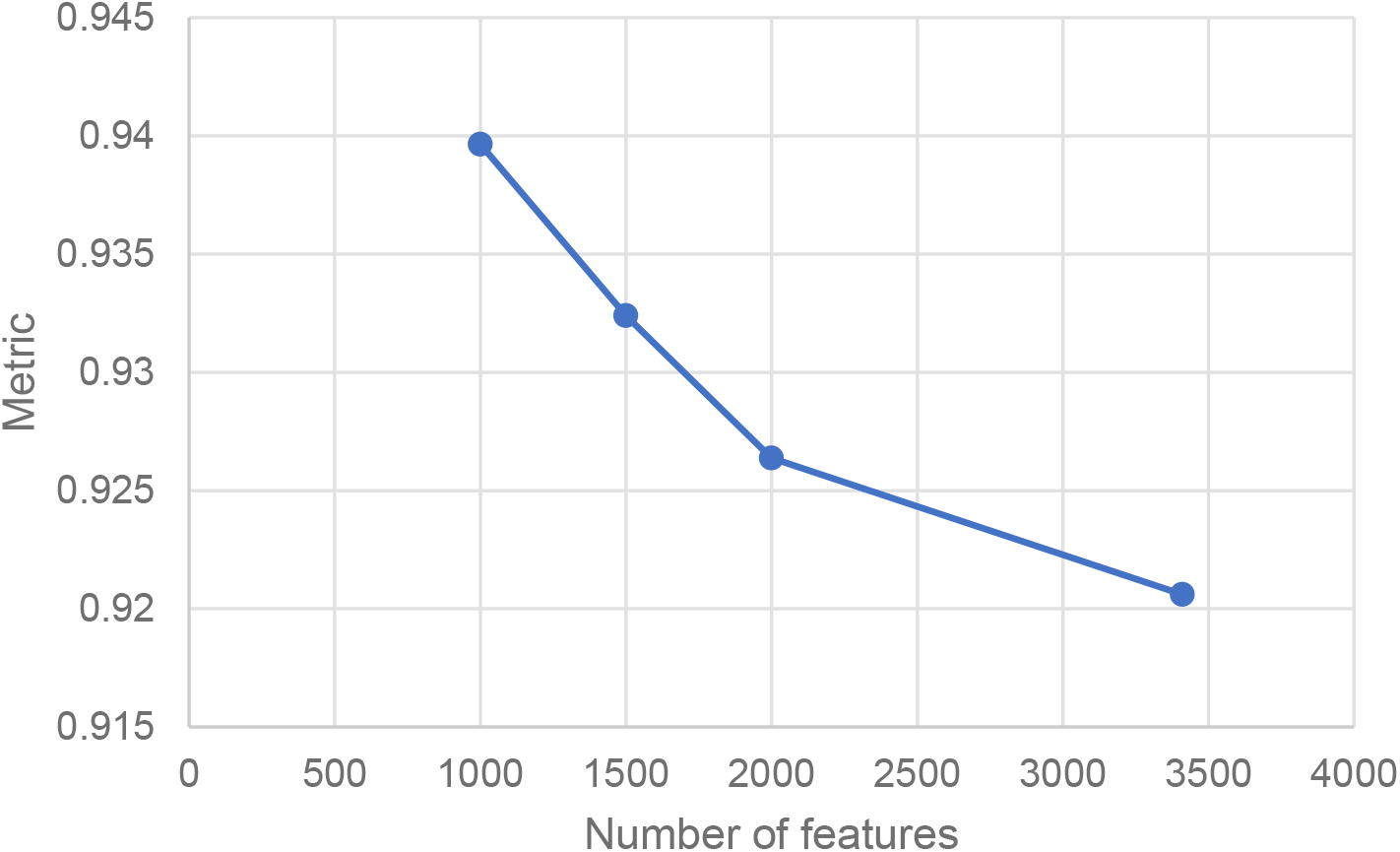
The relationship between the number of features and the value of metrics.

**Extended Data Fig. 2.**
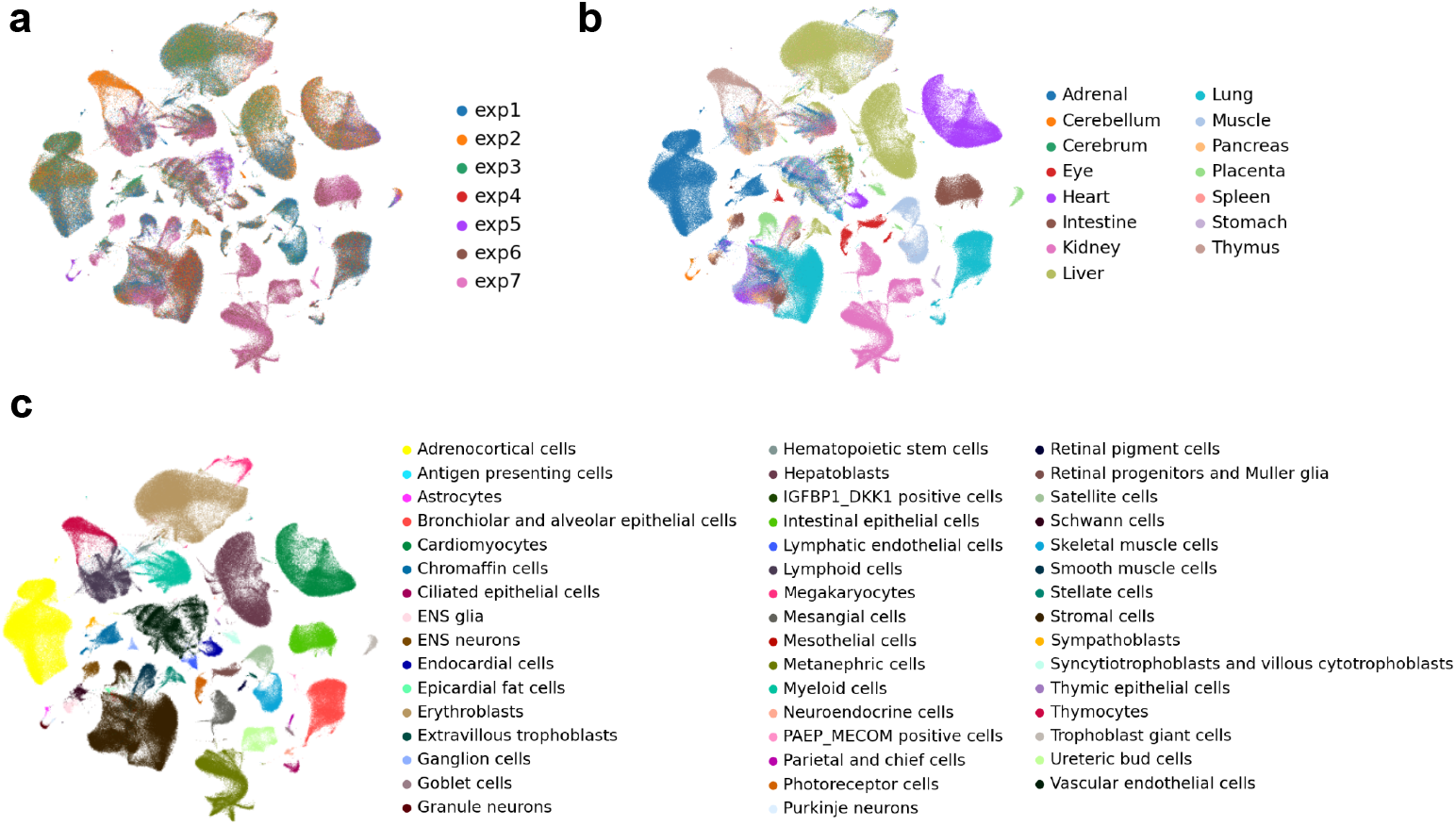
Visualization of cell embedding from the atlas-level dataset. (a) The UMAP colored by the batch label. (b) The UMAP colored by the organ label. (c) The UMAP colored by the cell type.

**Extended Data Fig. 3.**
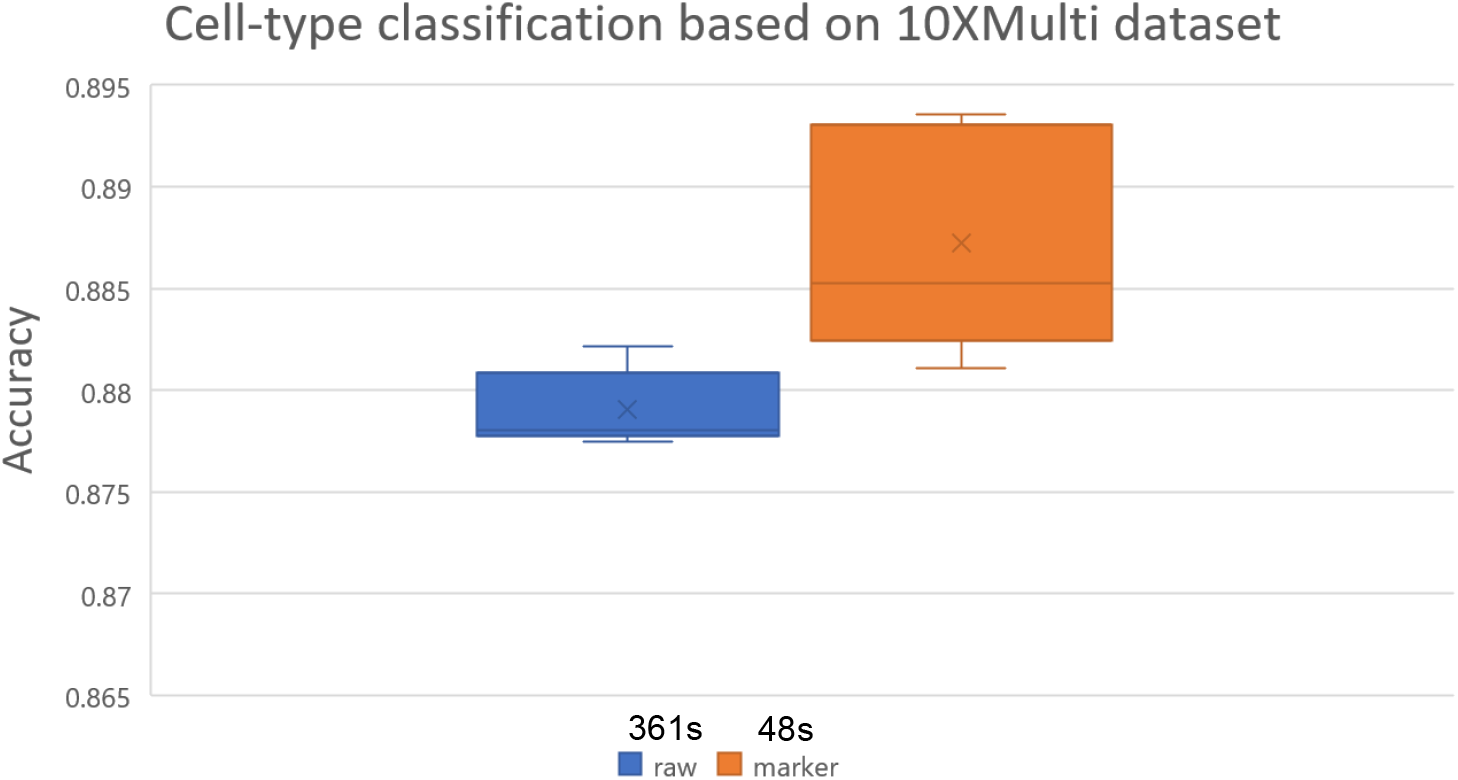
Comparisons of classification performances between all-gene case and selected-gene case. We display the boxplots based on five-fold cross validation. The accuracy and running time of classification is computed based on the pbmc neurips dataset.

**Extended Data Fig. 4.**
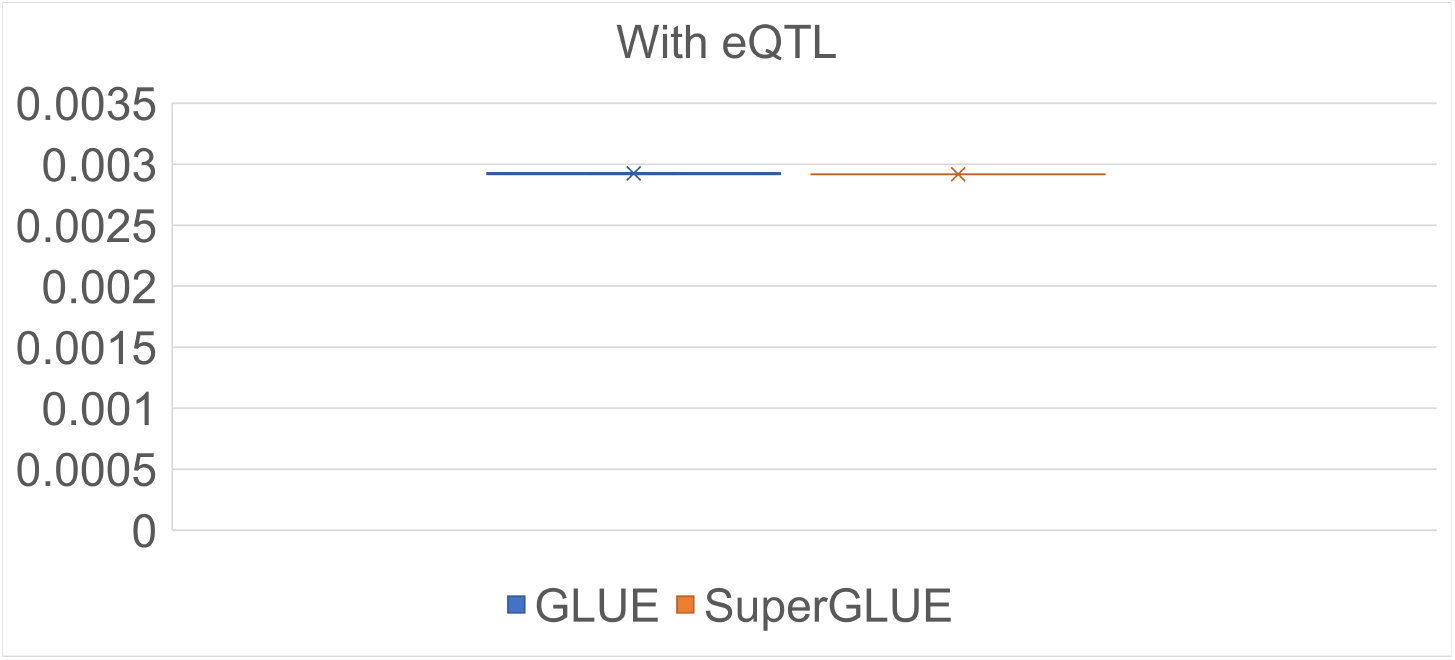
Comparisons of the overlap values between selected methods and eQTL ground truth.

**Extended Data Fig. 5.**
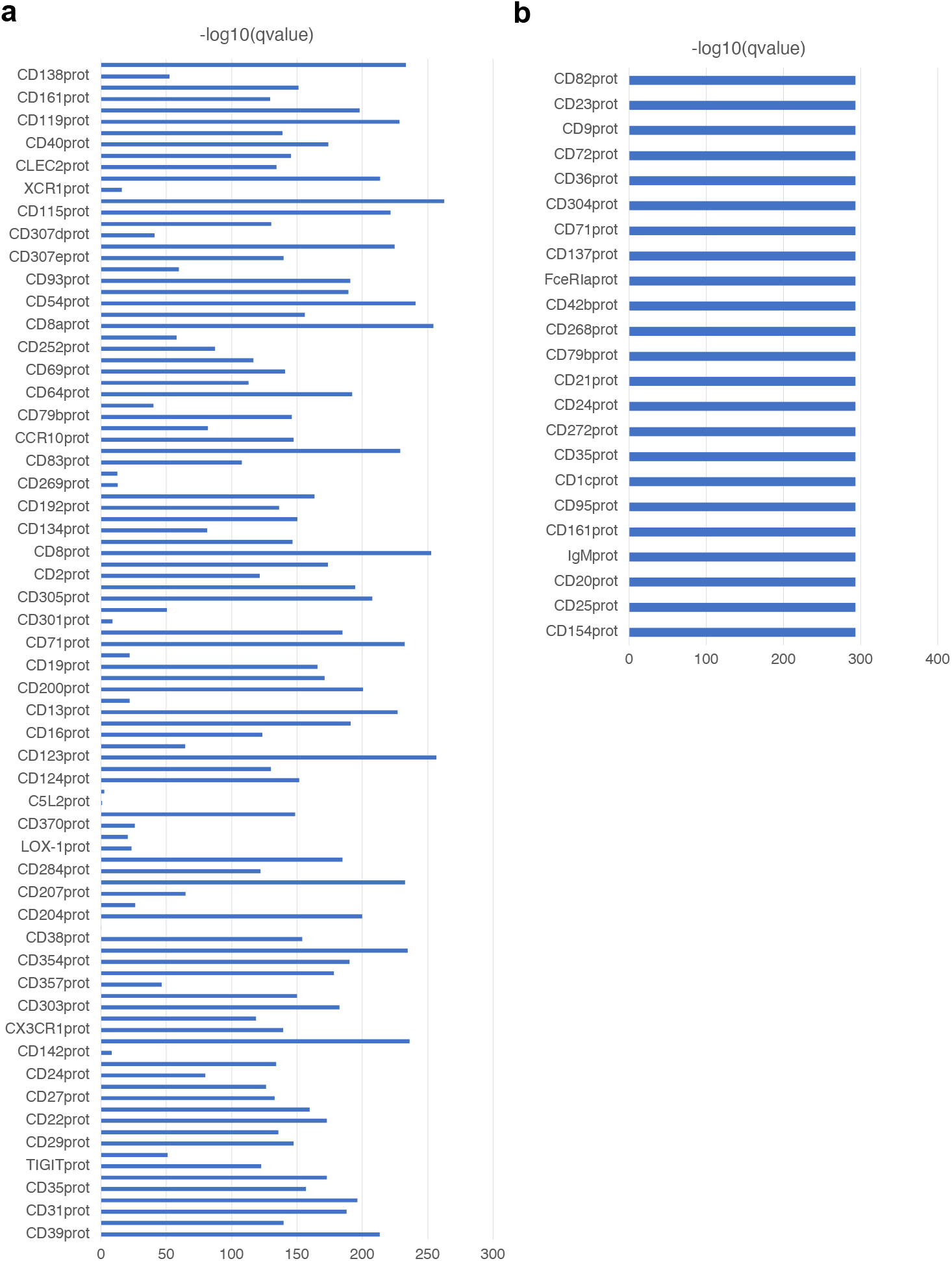
Testing statistics based on graph perturbation for two cite-seq datasets. (a) *−log*(*q − value*) of pbmc_seurat dataset. (b) *−log*(*q − value*) of pbmc competition dataset.

**Extended Data Fig. 6.**
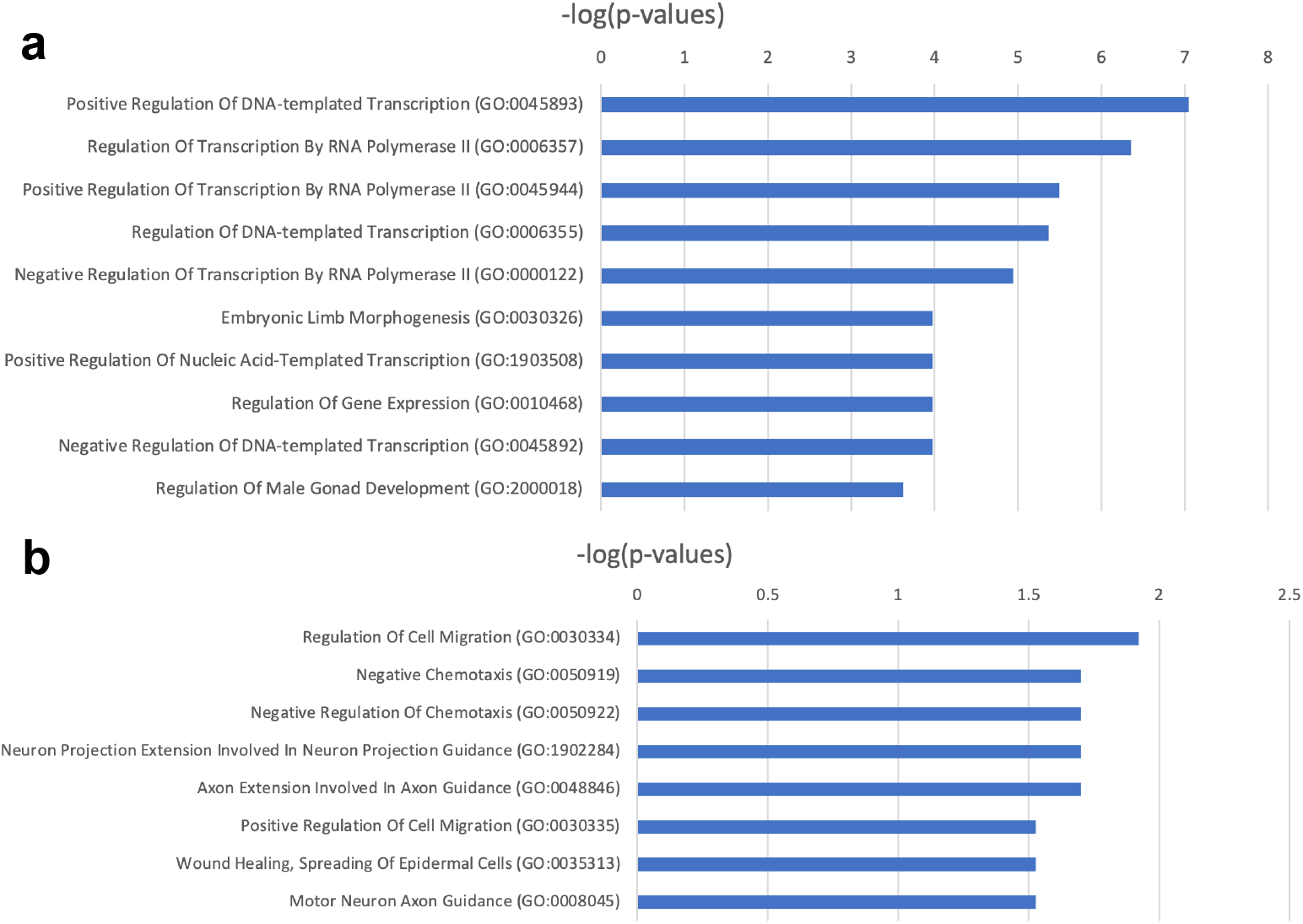
The comparison of GOEA results based on different gene sets. (a) The GOEA result of genes based the wild-type dataset. (b) The GOEA result of genes based the perturbed dataset.

**Extended Data Fig. 7.**
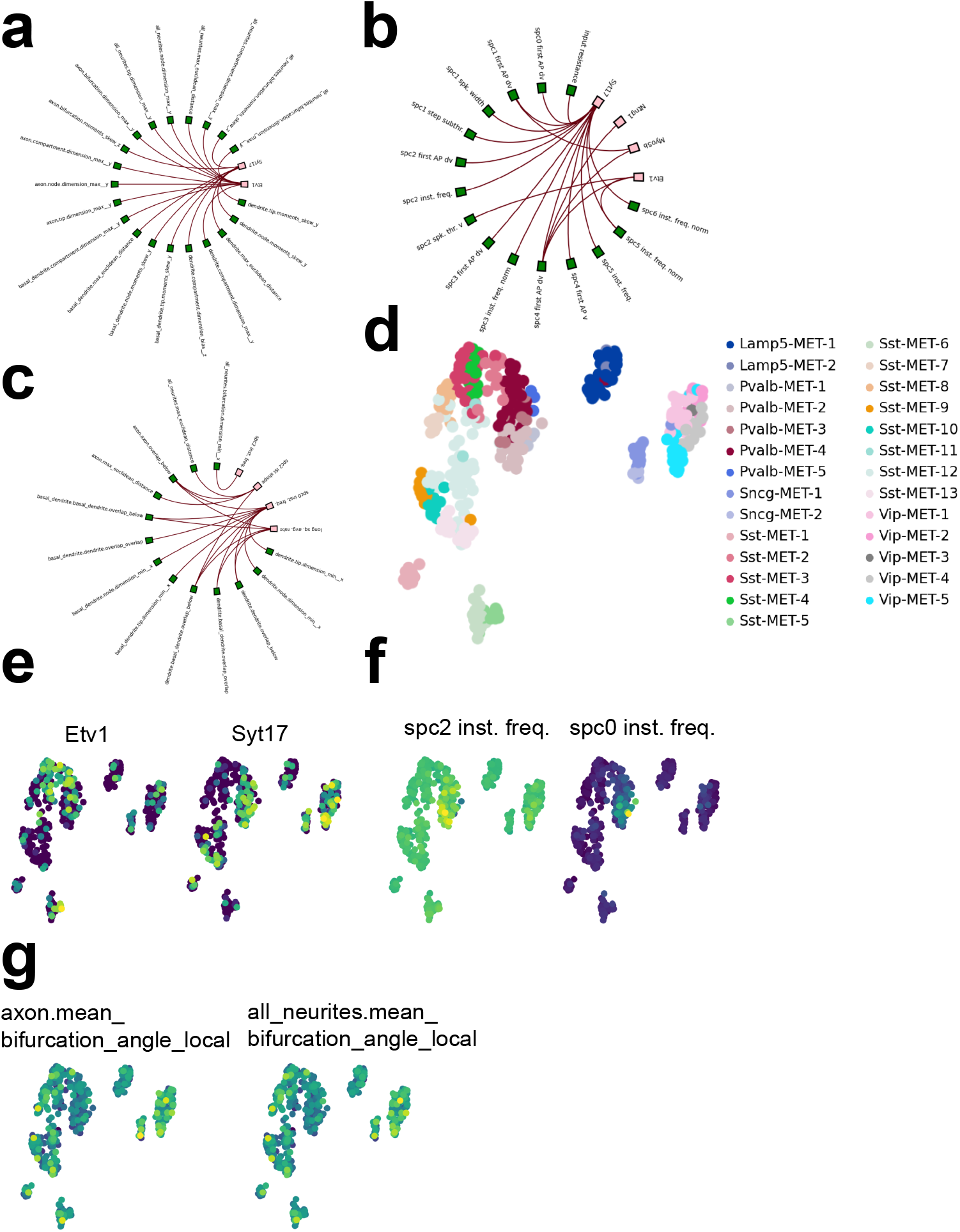
Analyses of the integration for the multi-sensing dataset. (a) The visualization of interactions between genes and morphology features. (b) The visualization of interactions between genes and electrophysiology features. (c) The visualization of interactions between morphology features and electrophysiology features. (d) The UMAP plot of cell embeddings colored by cell types. (e) The UMAP plot of cell embeddings colored by the expression levels of selected genes. (f) The UMAP plot of cell embeddings colored by the expression levels of morphology features. (g) The UMAP plot of cell embeddings colored by the expression levels of electrophysiology features.

**Extended Data Fig. 8.**
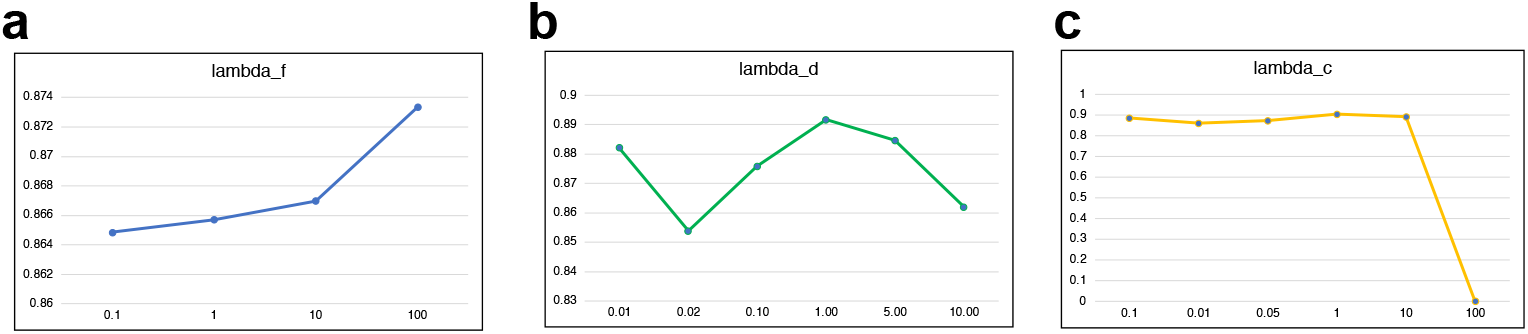
Results of hyper-parameter tuning. (a)-(c) correspond to hyper-parameters including *λ*_*f*_, *λ*_*d*_ and *λ*_*c*_.

## References

[1] Hwang, B., Lee, J.H., Bang, D.: Single-cell rna sequencing technologies and bioinformatics pipelines. Experimental & molecular medicine 50(8), 1–14 (2018)

[2] Zheng, G.X., Terry, J.M., Belgrader, P., Ryvkin, P., Bent, Z.W., Wilson, R., Ziraldo, S.B., Wheeler, T.D., McDermott, G.P., Zhu, J., et al.: Massively parallel digital transcriptional profiling of single cells. Nature communications 8(1), 1–12 (2017)

[3] Vandereyken, K., Sifrim, A., Thienpont, B., Voet, T.: Methods and applications for single-cell and spatial multi-omics. Nature Reviews Genetics 24(8), 494–515 (2023)

[4] Miao, Z., Humphreys, B.D., McMahon, A.P., Kim, J.: Multi-omics integration in the age of million single-cell data. Nature Reviews Nephrology 17(11), 710–724 (2021)

[5] Buenrostro, J.D., Wu, B., Litzenburger, U.M., Ruff, D., Gonzales, M.L., Snyder, M.P., Chang, H.Y., Greenleaf, W.J.: Single-cell chromatin accessibility reveals principles of regulatory variation. Nature 523(7561), 486–490 (2015)

[6] Stoeckius, M., Hafemeister, C., Stephenson, W., Houck-Loomis, B., Chattopadhyay, P.K., Swerdlow, H., Satija, R., Smibert, P.: Simultaneous epitope and transcriptome measurement in single cells. Nature methods 14(9), 865–868 (2017)

[7] Karemaker, I.D., Vermeulen, M.: Single-cell dna methylation profiling: technologies and biological applications. Trends in biotechnology 36(9), 952–965 (2018)

[8] Liu, Y., Yang, M., Deng, Y., Su, G., Enninful, A., Guo, C.C., Tebaldi, T., Zhang, D., Kim, D., Bai, Z., et al.: High-spatial-resolution multi-omics sequencing via deterministic barcoding in tissue. Cell 183(6), 1665–1681 (2020)

[9] Cadwell, C.R., Palasantza, A., Jiang, X., Berens, P., Deng, Q., Yilmaz, M., Reimer, J., Shen, S., Bethge, M., Tolias, K.F., et al.: Electrophysiological, transcriptomic and morphologic profiling of single neurons using patch-seq. Nature biotechnology 34(2), 199–203 (2016)

[10] Regev, A., Teichmann, S.A., Lander, E.S., Amit, I., Benoist, C., Birney, E., Bodenmiller, B., Campbell, P., Carninci, P., Clatworthy, M., et al.: The human cell atlas. elife 6, 27041 (2017)

[11] Bennett, D.A., Buchman, A.S., Boyle, P.A., Barnes, L.L., Wilson, R.S., Schneider, J.A.: Religious orders study and rush memory and aging project. Journal of Alzheimer’s disease 64(1), 161–189 (2018)

[12] Lee, M.Y., Kaestner, K.H., Li, M.: Benchmarking algorithms for joint integration of unpaired and paired single-cell rna-seq and atac-seq data. Genome Biology 24(1), 244 (2023)

[13] Xiao, C., Chen, Y., Meng, Q., Wei, L., Zhang, X.: Benchmarking multiomics integration algorithms across single-cell rna and atac data. Briefings in Bioinformatics 25(2), 095 (2024)

[14] Hu, Y., Wan, S., Luo, Y., Li, Y., Wu, T., Deng, W., Jiang, C., Jiang, S., Zhang, Y., Liu, N., et al.: Benchmarking algorithms for single-cell multi-omics prediction and integration. Nature Methods, 1–13 (2024)

[15] Hao, Y., Hao, S., Andersen-Nissen, E., Mauck, W.M., Zheng, S., Butler, A., Lee, M.J., Wilk, A.J., Darby, C., Zager, M., et al.: Integrated analysis of multimodal single-cell data. Cell 184(13), 3573–3587 (2021)

[16] Cao, Z.-J., Gao, G.: Multi-omics single-cell data integration and regulatory inference with graph-linked embedding. Nature Biotechnology 40(10), 1458–1466 (2022)

[17] Tang, X., Zhang, J., He, Y., Zhang, X., Lin, Z., Partarrieu, S., Hanna, E.B., Ren, Z., Shen, H., Yang, Y., et al.: Explainable multi-task learning for multi-modality biological data analysis. Nature communications 14(1), 2546 (2023)

[18] Gayoso, A., Steier, Z., Lopez, R., Regier, J., Nazor, K.L., Streets, A., Yosef, N.: Joint probabilistic modeling of single-cell multi-omic data with totalvi. Nature methods 18(3), 272–282 (2021)

[19] Ashuach, T., Gabitto, M.I., Koodli, R.V., Saldi, G.-A., Jordan, M.I., Yosef, N.: Multivi: deep generative model for the integration of multimodal data. Nature Methods 20(8), 1222–1231 (2023)

[20] Jain, M.S., Polanski, K., Conde, C.D., Chen, X., Park, J., Mamanova, L., Knights, A., Botting, R.A., Stephenson, E., Haniffa, M., et al.: Multimap: dimensionality reduction and integration of multimodal data. Genome biology 22, 1–26 (2021)

[21] Long, Y., Ang, K.S., Sethi, R., Liao, S., Heng, Y., Olst, L., Ye, S., Zhong, C., Xu, H., Zhang, D., et al.: Deciphering spatial domains from spatial multi-omics with spatialglue. Nature Methods, 1–10 (2024)

[22] Davidson, E., Levin, M.: Gene regulatory networks. Proceedings of the National Academy of Sciences 102(14), 4935–4935 (2005)

[23] Lundberg, S.M., Lee, S.-I.: A unified approach to interpreting model predictions. Advances in neural information processing systems 30 (2017)

[24] Kobak, D., Linderman, G.C.: Initialization is critical for preserving global data structure in both t-sne and umap. Nature biotechnology 39(2), 156–157 (2021)

[25] Kullback, S., Leibler, R.A.: On information and sufficiency. The annals of mathematical statistics 22(1), 79–86 (1951)

[26] Kipf, T.N., Welling, M.: Semi-Supervised Classification with Graph Convolutional Networks. OpenReview.net (2017). https://openreview.net/forum?id=SJU4ayYgl

[27] Liu, T., Wang, Y., Ying, R., Zhao, H.: Muse-gnn: Learning unified gene representation from multimodal biological graph data. Advances in Neural Information Processing Systems 36 (2024)

[28] Zügner, D., Borchert, O., Akbarnejad, A., Günnemann, S.: Adversarial attacks on graph neural networks: Perturbations and their patterns. ACM Transactions on Knowledge Discovery from Data (TKDD) 14(5), 1–31 (2020)

[29] Liu, X., Zhang, Y., Wu, M., Yan, M., He, K., Yan, W., Pan, S., Ye, X., Fan, D.: Revisiting edge perturbation for graph neural network in graph data augmentation and attack. arXiv preprint 2403.07943 (2024)

[30] Kobak, D., Berens, P.: The art of using t-sne for single-cell transcriptomics. Nature communications 10(1), 5416 (2019)

[31] Narayan, A., Berger, B., Cho, H.: Assessing single-cell transcriptomic variability through density-preserving data visualization. Nature biotechnology 39(6), 765– 774 (2021)

[32] Xu, Y., Zang, Z., Xia, J., Tan, C., Geng, Y., Li, S.Z.: Structure-preserving visualization for single-cell rna-seq profiles using deep manifold transformation with batch-correction. Communications Biology 6(1), 369 (2023)

[33] Luecken, M.D., Büttner, M., Chaichoompu, K., Danese, A., Interlandi, M., Müller, M.F., Strobl, D.C., Zappia, L., Dugas, M., Colomé-Tatché, M., et al.: Benchmarking atlas-level data integration in single-cell genomics. Nature methods 19(1), 41–50 (2022)

[34] Chari, T., Pachter, L.: The specious art of single-cell genomics. PLOS Computational Biology 19(8), 1011288 (2023)

[35] Lause, J., Kobak, D., Berens, P.: The art of seeing the elephant in the room: 2d embeddings of single-cell data do make sense. bioRxiv, 2024–03 (2024)

[36] Wolf, F.A., Hamey, F.K., Plass, M., Solana, J., Dahlin, J.S., Göttgens, B., Rajewsky, N., Simon, L., Theis, F.J.: Paga: graph abstraction reconciles clustering with trajectory inference through a topology preserving map of single cells. Genome biology 20, 1–9 (2019)

[37] Saelens, W., Cannoodt, R., Todorov, H., Saeys, Y.: A comparison of single-cell trajectory inference methods. Nature biotechnology 37(5), 547–554 (2019)

[38] Liu, T., Long, W., Cao, Z., Wang, Y., He, C.H., Zhang, L., Strittmatter, S.M., Zhao, H.: Cosgenegate selects multi-functional and credible biomarkers for single-cell analysis. bioRxiv, 2024–05 (2024)

[39] Cannoodt, R., Saelens, W., Deconinck, L., Saeys, Y.: Spearheading future omics analyses using dyngen, a multi-modal simulator of single cells. Nature Communications 12(1), 3942 (2021)

[40] Abdi, H., Williams, L.J.: Principal component analysis. Wiley interdisciplinary reviews: computational statistics 2(4), 433–459 (2010)

[41] Kingma, D.P., Welling, M.: Auto-encoding variational {Bayes}. In: Int. Conf. on Learning Representations

[42] Rumelhart, D.E., Hinton, G.E., Williams, R.J.: Learning internal representations by error propagation, parallel distributed processing, explorations in the microstructure of cognition, ed. de rumelhart and j. mcclelland. vol. 1. 1986. Biometrika 71, 599–607 (1986)

[43] Liu, T., Li, K., Wang, Y., Li, H., Zhao, H.: Evaluating the utilities of foundation models in single-cell data analysis. bioRxiv, 2023–09 (2023)

[44] Letsche, T.A., Berry, M.W.: Large-scale information retrieval with latent semantic indexing. Information sciences 100(1-4), 105–137 (1997)

[45] Granja, J.M., Corces, M.R., Pierce, S.E., Bagdatli, S.T., Choudhry, H., Chang, H.Y., Greenleaf, W.J.: Archr is a scalable software package for integrative single-cell chromatin accessibility analysis. Nature genetics 53(3), 403–411 (2021)

[46] Ma, S., Zhang, B., LaFave, L.M., Earl, A.S., Chiang, Z., Hu, Y., Ding, J., Brack, A., Kartha, V.K., Tay, T., et al.: Chromatin potential identified by shared single-cell profiling of rna and chromatin. Cell 183(4), 1103–1116 (2020)

[47] Chen, S., Lake, B.B., Zhang, K.: High-throughput sequencing of the transcriptome and chromatin accessibility in the same cell. Nature biotechnology 37(12), 1452–1457 (2019)

[48] Luecken, M.D., Burkhardt, D.B., Cannoodt, R., Lance, C., Agrawal, A., Aliee, H., Chen, A.T., Deconinck, L., Detweiler, A.M., Granados, A.A., et al.: A sandbox for prediction and integration of dna, rna, and proteins in single cells. In: Thirty-fifth Conference on Neural Information Processing Systems Datasets and Benchmarks Track (Round 2) (2021)

[49] Dou, J., Liang, S., Mohanty, V., Miao, Q., Huang, Y., Liang, Q., Cheng, X., Kim, S., Choi, J., Li, Y., et al.: Bi-order multimodal integration of single-cell data. Genome biology 23(1), 112 (2022)

[50] Stuart, T., Butler, A., Hoffman, P., Hafemeister, C., Papalexi, E., Mauck, W.M., Hao, Y., Stoeckius, M., Smibert, P., Satija, R.: Comprehensive integration of single-cell data. cell 177(7), 1888–1902 (2019)

[51] Arunachalam, P.S., Wimmers, F., Mok, C.K.P., Perera, R.A., Scott, M., Hagan, T., Sigal, N., Feng, Y., Bristow, L., Tak-Yin Tsang, O., et al.: Systems biological assessment of immunity to mild versus severe covid-19 infection in humans. Science 369(6508), 1210–1220 (2020)

[52] Luo, C., Keown, C.L., Kurihara, L., Zhou, J., He, Y., Li, J., Castanon, R., Lucero, J., Nery, J.R., Sandoval, J.P., et al.: Single-cell methylomes identify neuronal subtypes and regulatory elements in mammalian cortex. Science 357(6351), 600–604 (2017)

[53] Saunders, A., Macosko, E.Z., Wysoker, A., Goldman, M., Krienen, F.M., Rivera, H., Bien, E., Baum, M., Bortolin, L., Wang, S., et al.: Molecular diversity and specializations among the cells of the adult mouse brain. Cell 174(4), 1015–1030 (2018)

[54] Gouwens, N.W., Sorensen, S.A., Baftizadeh, F., Budzillo, A., Lee, B.R., Jarsky, T., Alfiler, L., Baker, K., Barkan, E., Berry, K., et al.: Integrated morphoelectric and transcriptomic classification of cortical gabaergic cells. Cell 183(4), 935–953 (2020)

[55] Finn, E.S., Huber, L., Jangraw, D.C., Molfese, P.J., Bandettini, P.A.: “Layer-dependent Activity in Human Prefrontal Cortex During Working Memory”. 10.18112/openneuro.ds002076.v1.0.1

[56] Xiong, L., Chen, T., Kellis, M.: scclip: Multi-modal single-cell contrastive learning integration pre-training. In: NeurIPS 2023 AI for Science Workshop (2023)

[57] Cui, H., Wang, C., Maan, H., Pang, K., Luo, F., Duan, N., Wang, B.: scgpt: toward building a foundation model for single-cell multi-omics using generative ai. Nature Methods, 1–11 (2024)

[58] Mercatelli, D., Scalambra, L., Triboli, L., Ray, F., Giorgi, F.M.: Gene regulatory network inference resources: A practical overview. Biochimica et Biophysica Acta (BBA)-Gene Regulatory Mechanisms 1863(6), 194430 (2020)

[59] Navlakha, S., Bar-Joseph, Z.: Algorithms in nature: the convergence of systems biology and computational thinking. Molecular systems biology 7(1), 546 (2011)

[60] LeCun, Y., Bengio, Y., Hinton, G.: Deep learning. nature 521(7553), 436–444 (2015)

[61] Webb, S., et al.: Deep learning for biology. Nature 554(7693), 555–557 (2018)

[62] Castelvecchi, D.: Can we open the black box of ai? Nature News 538(7623), 20 (2016)

[63] Pullin, J.M., McCarthy, D.J.: A comparison of marker gene selection methods for single-cell rna sequencing data. Genome Biology 25(1), 56 (2024)

[64] Wallace, D.C.: Mitochondrial dna mutations in disease and aging. Environmental and molecular mutagenesis 51(5), 440–450 (2010)

[65] Malier, M., Gharzeddine, K., Laverriere, M.-H., Marsili, S., Thomas, F., Court, M., Decaens, T., Roth, G., Millet, A.: Hypoxia drives dihydropyrimidine dehydrogenase expression in macrophages and confers chemoresistance in colorectal cancer. Cancer Research 81(23), 5963–5976 (2021)

[66] Xie, L., Zhang, S., Huang, L., Peng, Z., Lu, H., He, Q., Chen, R., Hu, L., Wang, B., Sun, B., et al.: Single-cell rna sequencing of peripheral blood reveals that monocytes with high cathepsin s expression aggravate cerebral ischemia– reperfusion injury. Brain, Behavior, and Immunity 107, 330–344 (2023)

[67] Javierre, B.M., Burren, O.S., Wilder, S.P., Kreuzhuber, R., Hill, S.M., Sewitz, S., Cairns, J., Wingett, S.W., Várnai, C., Thiecke, M.J., et al.: Lineage-specific genome architecture links enhancers and non-coding disease variants to target gene promoters. Cell 167(5), 1369–1384 (2016)

[68] Aguet François 1 Brown Andrew A. 2 3 4 Castel Stephane E. 5 6 Davis Joe R. 8 He Yuan 9 Jo Brian 10 Mohammadi Pejman 5 6 Park YoSon 11 Parsana Princy 12 Segré Ayellet V. 1 Strober Benjamin J. 9 Zappala Zachary 7 8, G.C.L., Addington Anjene 15 Guan Ping 16 Koester Susan 15 Little A. Roger 17 Lockhart Nicole C. 18 Moore Helen M. 16 Rao Abhi 16 Struewing Jeffery P. 19 Volpi Simona 19, N., 16, P.S.L..B.M.E..B.P.A., 137, N.C.F.N.C.R., et al.: Genetic effects on gene expression across human tissues. Nature 550(7675), 204–213 (2017)

[69] Aibar, S., González-Blas, C.B., Moerman, T., Huynh-Thu, V.A., Imrichova, H., Hulselmans, G., Rambow, F., Marine, J.-C., Geurts, P., Aerts, J., et al.: Scenic: single-cell regulatory network inference and clustering. Nature methods 14(11), 1083–1086 (2017)

[70] Bravo González-Blas, C., De Winter, S., Hulselmans, G., Hecker, N., Matetovici, I., Christiaens, V., Poovathingal, S., Wouters, J., Aibar, S., Aerts, S.: Scenic+: single-cell multiomic inference of enhancers and gene regulatory networks. Nature methods 20(9), 1355–1367 (2023)

[71] Li, M.M., Huang, Y., Sumathipala, M., Liang, M.Q., Valdeolivas, A., Ananthakrishnan, A.N., Liao, K., Marbach, D., Zitnik, M.: Contextual ai models for single-cell protein biology. Nature Methods, 1–12 (2024)

[72] Gielen, P.R., Schulte, B.M., Kers-Rebel, E.D., Verrijp, K., Petersen-Baltussen, H.M., Ter Laan, M., Wesseling, P., Adema, G.J.: Increase in both cd14-positive and cd15-positive myeloid-derived suppressor cell subpopulations in the blood of patients with glioma but predominance of cd15-positive myeloid-derived suppressor cells in glioma tissue. Journal of neuropathology & experimental neurology 74(5), 390–400 (2015)

[73] Borvak, J., Chou, C.-S., Bell, K., Van Dyke, G., Zola, H., Ramilo, O., Vitetta, E.S.: Expression of cd25 defines peripheral blood mononuclear cells with productive versus latent hiv infection. Journal of immunology (Baltimore, Md.: 1950) 155(6), 3196–3204 (1995)

[74] Liu, X., Li, Q., Zhou, Y., He, X., Fang, J., Lu, H., Wang, X., Wang, D., Ma, D., Cheng, B., et al.: Dysfunctional role of elevated tigit expression on t cells in oral squamous cell carcinoma patients. Oral Diseases 27(7), 1667–1677 (2021)

[75] Teillaud, C., Galon, J., Zilber, M.-T., Mazieres, N., Spagnoli, R., Kurrle, R., Fridman, W.H., Sautes, C.: Soluble cd16 binds peripheral blood mononuclear cells and inhibits pokeweed-mitogen-induced responses (1993)

[76] Schiller, A., Zhang, T., Li, R., Duechting, A., Sundararaman, S., Przybyla, A., Kuerten, S., Lehmann, P.V.: A positive control for detection of functional cd4 t cells in pbmc: the cpi pool. Cells 6(4), 47 (2017)

[77] Vudattu, N.K., Magalhaes, I., Schmidt, M., Seyfert-Margolis, V., Maeurer, M.J.: Reduced numbers of il-7 receptor (cd127) expressing immune cells and il-7-signaling defects in peripheral blood from patients with breast cancer. International journal of cancer 121(7), 1512–1519 (2007)

[78] Sconocchia, G., Keyvanfar, K., El Ouriaghli, F., Grube, M., Rezvani, K., Fujiwara, H., McCoy, J., Hensel, N., Barrett, A.: Phenotype and function of a cd56+ peripheral blood monocyte. Leukemia 19(1), 69–76 (2005)

[79] De Oliveira, S.N., Wang, J., Ryan, C., Morrison, S.L., Kohn, D.B., Hollis, R.P.: A cd19/fc fusion protein for detection of anti-cd19 chimeric antigen receptors. Journal of translational medicine 11, 1–9 (2013)

[80] Kern, F., Faulhaber, N., Frömmel, C., Khatamzas, E., Prösch, S., Schönemann, 27 C., Kretzschmar, I., Volkmer-Engert, R., Volk, H.-D., Reinke, P.: Analysis of cd8 t cell reactivity to cytomegalovirus using protein-spanning pools of overlapping pentadecapeptides. European journal of immunology 30(6), 1676–1682 (2000)

[81] Landmann, R., Knopf, H.-P., Link, S., Sansano, S., Schumann, R., Zimmerli, W.: Human monocyte cd14 is upregulated by lipopolysaccharide. Infection and immunity 64(5), 1762–1769 (1996)

[82] Hecker, M., Lambeck, S., Toepfer, S., Van Someren, E., Guthke, R.: Gene regulatory network inference: data integration in dynamic models—a review. Biosystems 96(1), 86–103 (2009)

[83] Argelaguet, R., Lohoff, T., Li, J.G., Nakhuda, A., Drage, D., Krueger, F., Velten, L., Clark, S.J., Reik, W.: Decoding gene regulation in the mouse embryo using single-cell multi-omics. BioRxiv, 2022–06 (2022)

[84] Weaver, J.C., Chizmadzhev, Y.A.: Theory of electroporation: a review. Bioelectrochemistry and bioenergetics 41(2), 135–160 (1996)

[85] Tanaka, M., Nakamura, T.: Targeting epigenetic aberrations of sarcoma in crispr era. Genes, Chromosomes and Cancer 62(9), 510–525 (2023)

[86] Overeem, A.W., Chang, Y.W., Moustakas, I., Roelse, C.M., Hillenius, S., Van Der Helm, T., Van Der Schrier, V.F., Gonçalves, M.A., Mei, H., Freund, C., et al.: Efficient and scalable generation of primordial germ cells in 2d culture using basement membrane extract overlay. Cell Reports Methods 3(6) (2023)

[87] Deng, C., Zhang, Z., Xu, F., Xu, J., Ren, Z., Godoy-Parejo, C., Xiao, X., Liu, W., Zhou, Z., Chen, G.: Thyroid hormone enhances stem cell maintenance and promotes lineage-specific differentiation in human embryonic stem cells. Stem Cell Research & Therapy 13(1), 120 (2022)

[88] Thomas, P.D., Ebert, D., Muruganujan, A., Mushayahama, T., Albou, L.-P., Mi, H.: Panther: Making genome-scale phylogenetics accessible to all. Protein Science 31(1), 8–22 (2022)

[89] Yang, Z., Ding, M., Zhou, C., Yang, H., Zhou, J., Tang, J.: Understanding negative sampling in graph representation learning. In: Proceedings of the 26th ACM SIGKDD International Conference on Knowledge Discovery & Data Mining, pp. 1666–1676 (2020)

[90] Goodfellow, I., Pouget-Abadie, J., Mirza, M., Xu, B., Warde-Farley, D., Ozair, S., Courville, A., Bengio, Y.: Generative adversarial networks. Communications of the ACM 63(11), 139–144 (2020)

[91] Seabold, S., Perktold, J.: Statsmodels: econometric and statistical modeling with python. SciPy 7, 1 (2010)

[92] Davis, C.A., Hitz, B.C., Sloan, C.A., Chan, E.T., Davidson, J.M., Gabdank, I., Hilton, J.A., Jain, K., Baymuradov, U.K., Narayanan, A.K., et al.: The encyclopedia of dna elements (encode): data portal update. Nucleic acids research 46(D1), 794–801 (2018)

[93] Hagberg, A., Swart, P. S, Chult, D.: Exploring network structure, dynamics, and function using networkx. Technical report, Los Alamos National Lab.(LANL), Los Alamos, NM (United States) (2008)

[94] Gu, N., Vervaeke, K., Storm, J.F.: Bk potassium channels facilitate high-frequency firing and cause early spike frequency adaptation in rat ca1 hippocampal pyramidal cells. The Journal of physiology 580(3), 859–882 (2007)

[95] Yan, J., Aldrich, R.W.: Bk potassium channel modulation by leucine-rich repeat-containing proteins. Proceedings of the National Academy of Sciences 109(20), 7917–7922 (2012)

[96] Nooij, J.C., Doobar, S., Jessell, T.M.: Etv1 inactivation reveals proprioceptor subclasses that reflect the level of nt3 expression in muscle targets. Neuron 77(6), 1055–1068 (2013)

[97] Ruhl, D.A., Bomba-Warczak, E., Watson, E.T., Bradberry, M.M., Peterson, T.A., Basu, T., Frelka, A., Evans, C.S., Briguglio, J.S., Basta, T., et al.: Synaptotagmin 17 controls neurite outgrowth and synaptic physiology via distinct cellular pathways. Nature communications 10(1), 3532 (2019)

[98] Kaji, I., Roland, J.T., Rathan-Kumar, S., Engevik, A.C., Burman, A., Goldstein, A.E., Watanabe, M., Goldenring, J.R.: Cell differentiation is disrupted by myo5b loss through wnt/notch imbalance. JCI insight 6(16) (2021)

[99] Yaguchi, K., Nishimura-Akiyoshi, S., Kuroki, S., Onodera, T., Itohara, S.: Identification of transcriptional regulatory elements for ntng1 and ntng2 genes in mice. Molecular brain 7, 1–16 (2014)

[100] Kitajima, H., Kurths, J.: Bifurcation in neuronal networks with hub structure. Physica A: Statistical Mechanics and its Applications 388(20), 4499–4508 (2009)

